# Systems-level feedback loops maintain gene expression homeostasis following RNA polymerase II dosage perturbation

**DOI:** 10.1101/2025.09.18.677100

**Authors:** Anouk M. Olthof, Melvin J. Noe Gonzalez, Nanna Bach Poulsen, Nicolas Nieto Moreno, Theodora Bautu, Simona Cugusi, Jesper Q. Svejstrup

**Affiliations:** Center for Gene Expression, Department of Cellular and Molecular Medicine, Panum Institute, Blegdamsvej 3B, University of Copenhagen, 2200 Copenhagen N, Denmark; The Francis Crick Institute, 1 Midland Road, London NW1 1AT, UK

## Abstract

Transcription is regulated by sequence-specific transcription factors and enzymes allowing access to genes in chromatin. However, recent data indicate that the abundance of RNA polymerase II (RNAPII) itself may under certain circumstances represent an additional, crucial determinant of transcription regulation. Here, we used the dTAG system to titrate the cellular dosage of human RPB1, the largest RNAPII subunit, to more generally assess the importance of RNAPII levels. Unexpectedly, cells are extremely sensitive to RPB1 dosage, with a mere 30% reduction sufficient to perturb cell proliferation, cell cycle progression, and global transcription. Importantly, alterations in RPB1 abundance trigger hierarchical gene expression changes that are highly organized rather than stochastic. Using a combination of sequencing and proteomic approaches, we uncover the existence of multiple feedback loops between transcriptional initiation, promotor-proximal pause release, transcript elongation, splicing, and mRNA half-life, which together establish RNAPII abundance as a crucial systems-level regulator of transcriptional homeostasis.

## Introduction

In higher eukaryotes, RNA polymerase II (RNAPII) transcription is fundamentally shaped by chromatin structure and the binding of DNA sequence-specific transcription factors in the vicinity of gene promoters or at remote enhancer elements. These transcription factors determine which genes are active^1^, while chromatin modifications and remodeling establish a permissive environment for transcription^2–4^. In this prevalent view of transcription regulation, the transcription factor make-up of a cell alone ensures that genes are expressed at the right time, place, and level, which in turn is critical for development, differentiation, and the response to internal and external regulatory signals.

To meet the demands of cellular, tissue, and organismal specification, the correct expression and concentration of these transcription factors is tightly regulated^1,5^. Indeed, when a transcription factor is present at low concentrations, its binding to promoters becomes probabilistic^6,7^. This results in stochastic gene expression, which can have a negative impact on the overall fitness of a cell when it is uncontrolled^8,9^. Other evidence that cells are sensitive to changes in transcription factor levels stems from human genetics. Most Mendelian diseases are inherited in a recessive manner^10,11^, which implies that loss of one of the two copies of a gene often has little consequence and thus that a two-fold change in their expression may be tolerated. In contrast, more than two-thirds of all disease-causing mutations in genes encoding transcription factors are heterozygous^10,12,13^. In other words, half-normal levels of DNA binding transcription factors often disrupt normal cell function.

Despite its evident importance for transcription, RNAPII lacks the ability to faithfully initiate transcription by itself. Instead, general transcription factors and co-activators such Mediator are required to allow the formation of a pre-initiation complex and enable transcription initiation^14,15^, while additional pausing-, elongation-, and termination factors are required to support its further progress through the transcription cycle^16–18^. In the framework of transcription factor-driven gene regulation, the role of RNAPII itself is often overlooked or considered self-evident: it is the implicit assumption that RNAPII is recruited to genes from a plentiful nucleoplasmic pool to transcribe genes according to regulatory cues. Nevertheless, disease-causing mutations have been identified in subunits of RNAPII, and these are *de novo* heterozygous mutations in *POLR2A*, the gene encoding the main catalytic subunit RPB1^19,20^. Several of these mutations are thought to result in haploinsufficiency, raising the possibility that cellular regulation might be sensitive to changes in RPB1 levels.

Interestingly, because its abundance increases with cell size in yeast^21–23^, RNAPII has been suggested to be a limiting factor during biosynthetic scaling. This phenomenon, whereby the concentration of key proteins is maintained within a homeostatic range when cells increase in size, is well-documented, but the underlying mechanisms remain unclear. While RNAPII has recently been shown to be the limiting factor for the formation of the pre-initiation complex during cell size scaling in yeast^21^, the effect on mRNA production was not investigated, preventing a clear conclusion as to whether RNAPII levels were also limiting for transcription^24^. Indeed, findings in human cells suggested that changes in RNAPII levels are merely the consequence of a negative feedback loop between mRNA concentration and transcription^25^. It thus remains unclear whether human RNAPII is limiting for transcriptional activity and how sub-optimal levels affect basic gene expression parameters.

Evidence on the importance of regulating RPB1 levels stems from studies focused on the transcriptional response to UV-induced DNA damage. Specifically, it has been shown that regulation of the RNAPII pool size is essential for both the transcriptional shutdown and transcription restart that occur following UV-irradiation^26^. For obvious reasons, it has been difficult to more generally study how small changes in RNAPII levels affect gene expression in physiological conditions. Such experiments require controlled perturbation methods that are acute, in order to parse immediate versus late (adaptive) responses. Here, we generated a degron cell line for RPB1, which allows us to precisely titrate its levels using varying doses of dTAG^V^-1 degrader and determine the importance of RNAPII abundance. We found that a mere 30% reduction in RPB1 abundance is sufficient to drastically impair global transcription and affect the proliferation of cells. Furthermore, using a variety of sequencing techniques, we uncovered the presence of numerous feedback loops that buffer transcriptional output to changes in RPB1 dosage. Together, our findings show that cells are extremely sensitive to RPB1 dosage and that RNA polymerase II levels are tightly controlled by the cell in order to maintain the correct levels of steady-state gene expression.

## Results

### Precise modulation of RPB1 levels through the dTAG^V^-1 system

In order to study the role of RNAPII dosage in gene expression homeostasis, we generated a degron cell line, using the degradation tag (dTAG) system^27^ to facilitate removal of RPB1 upon addition of heterobifunctional dTAG molecules. Given previous observations that addition of a degradation tag at the C-terminus of RPB1 may result in degradation of only its C-terminal domain (CTD)^28,29^, we instead targeted the *POLR2A* gene region encoding the N-terminus of RPB1 (Figure 1A). Successful knock-in and expression of the degradation tag was confirmed by genotyping and the shift in RPB1 molecular weight by Western blot analysis (Supplementary Figure S1A-B). The introduction of the degradation tag did not alter RPB1 protein levels in these cells, and immunofluorescence with antibodies against the HA-tag and RPB1 showed that, as expected, treatment with 500nM dTAG^V^-1 resulted in degradation of RPB1 in all cells, rather than affecting just a subset of cells (Figure 1B-C).

**Figure 1.**
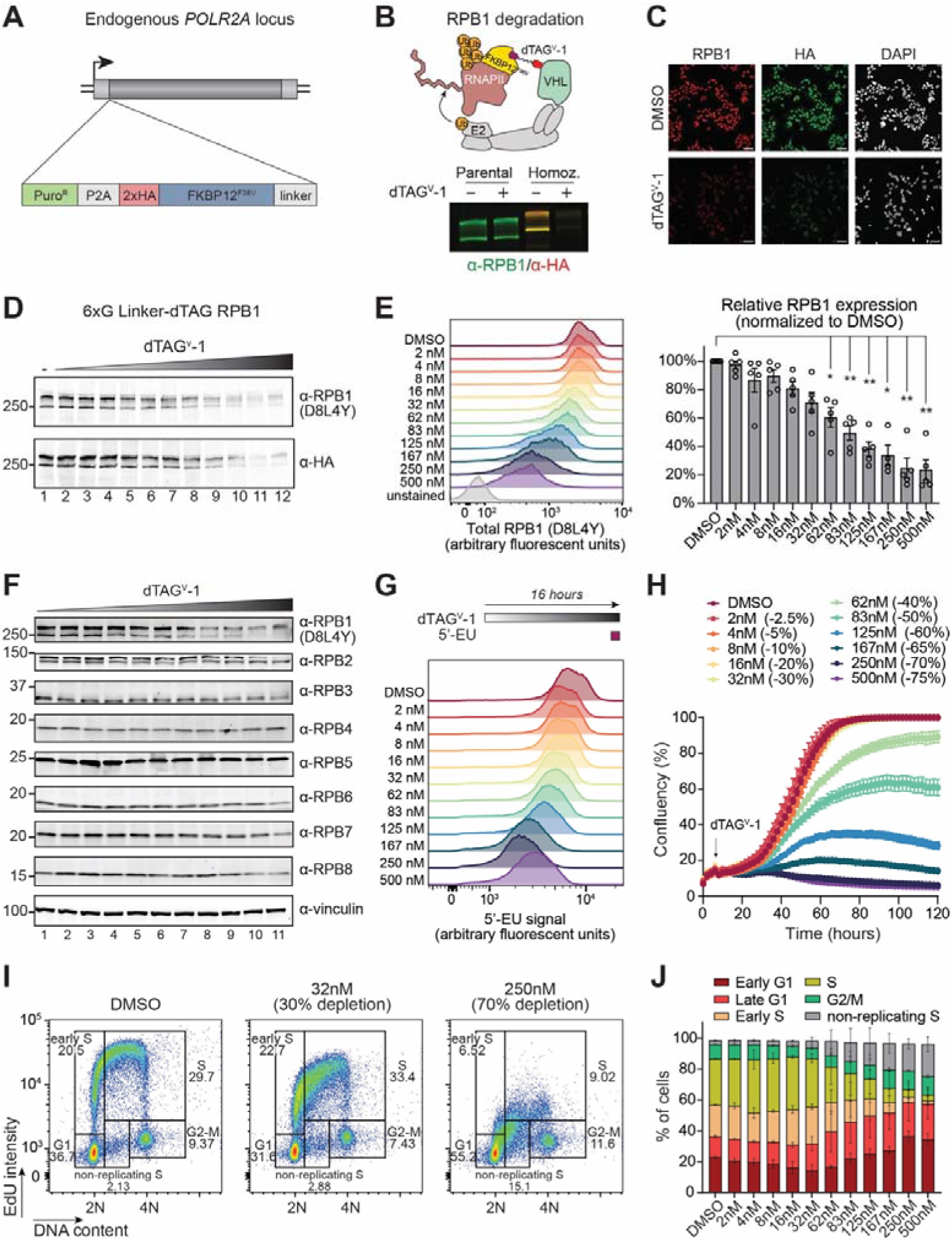
Precise modulation of RPB1 levels through the dTAG^V^-1 system. **(A)** Schematic of the degron construct knocked-in the endogenous *POLR2A* locus to generate RPB1 degron cell lines. **(B)** Schematic of dTAG^V^-1 mediated degradation of RPB1. **(C)** Immunofluorescence of RPB1 degron cells treated for 24 hours with 500nM dTAG^V^-1. RPB1 was detected using an antibody against the N-terminus of RPB1 and an anti-HA tag antibody **(D)** Western blot analysis of dose-dependent dTAG^V^-1-mediated degradation of RPB1 in the 6XG linker RPB1 degron cell line using an antibody against the N-terminus of RPB1 and an anti-HA tag antibody. dTAG^V^-1 concentrations are the same as in (E). **(E)** Flow cytometry analysis of total RPB1 levels following treatment with different doses of dTAG^V^-1. Quantification of the median RPB1 levels of at least four independent experiments are shown on the right. Bargraphs represent mean ±SEM. Statistical significance between DMSO and dTAG^V^-1 conditions was determined using a mixed-effects model, followed by Dunnett’s multiple comparisons test. *=P-value<0.05; **=P-value<0.01. **(F)** Western blot analysis of RNAPII subunit expression in the 6XG linker RPB1 degron cell line, following different dTAG^V^-1 concentrations. dTAG^V^-1 concentrations are the same as in (E). **(G)** Flow cytometry analysis of 5’-EU incorporation into nascent RNA following treatment with different doses of dTAG^V^-1. Cells were pulsed with 5’-EU for 1 hour prior to harvest. **(H)** Growth assay of 6XG linker RPB1 degron cells following treatment with different concentrations of dTAG^V^-1. Corresponding RPB1 depletion level is shown between brackets. Cell confluency was measured every 4hrs using Incucyte. Data is represented as the mean of three replicates ±SEM. **(I)** Flow cytometry analysis of cell cycle distributions. **(J)** Quantification of cell cycle distributions following treatment with different concentrations of dTAG^V^-1. Data is represented as the mean of two independent experiments ±SEM.

In an attempt to titrate RPB1 levels, we first expressed an RPB1 transgene under the control of a doxycycline-regulated hybrid CMV/TetO2 promoter in the RPB1 degron cell line (Supplementary Figure S1C). As expected, the addition of doxycycline successfully induced expression of the RPB1 transgene. However, titrating down the concentration of doxycycline did not result in a consistent dose-dependent reduction of RPB1 levels (Supplementary Figure S1D-F; and data not shown). We therefore decided to modulate the levels of RPB1 by simply titrating the concentration of dTAG^V^-1 instead.

Using multiple different concentrations of dTAG for 16 hours, we were able to obtain a range of RPB1 levels in distinct RPB1 degron clones (Figure 1D; Supplementary Figure S2A-B). Flow cytometry revealed a normal distribution for the expression of RPB1 not only in the DMSO-treated cells, but also in the different dTAG^V^-1-treated cells, albeit obviously at reduced levels (Figure 1E). Together, these experiments showed that different dTAG^V^-1 concentrations can be used to titrate RPB1 levels in a homogeneous and reproducible manner. Quantification of the flow cytometry data revealed that 500 nM dTAG^V^-1, the dose most frequently used in literature, after 16 hours resulted in a substantial 75% reduction of RPB1 levels, while 2nM dTAG^V^-1 reduced RPB1 levels by only 2.5% (Figure 1E; Supplementary Figure S2C). These effects were consistent across different experiments. The depletion of RPB1 affected the levels of other RNAPII subunits within this timeframe, but to a smaller extent (Figure 1F; Supplementary Figure S2D). To verify that the depletion of RPB1 indeed reduced transcription, we pulsed cells with 5’-ethynyluridine (5’-EU) for 1 hour to evaluate global transcription levels using flow cytometry (Figure 1G; Supplementary Figure S2E). Notably, 5’-EU is incorporated into all newly transcribed RNAs, including those synthesized by RNAPI and RNAPIII. 5’-EU pulsing combined with flow cytometry thus likely represents a major underestimate of the effect of RPB1 levels on transcription of protein-coding genes. Nevertheless, these results indicated that the dTAG system can be used to precisely modulate RPB1 levels and study the role of RNAPII dosage on gene expression.

### Depletion of RPB1 results in cell cycle arrest

To first determine whether cells are sensitive to RPB1 dosage, we treated with dTAG^V^-1 and followed cell growth using live imaging. This revealed that cells can only tolerate up to 30% reduction in the total RPB1 pool; any further reduction impacts cell growth (Figure 1H). We note that a 30% reduction is barely noticeable by a typical analysis approach such a Western blotting, so this provided a first indication that cells do not carry a great excess of RNAPII. Depletion of RPB1 resulted in activation of apoptotic signaling (Supplementary Figure S3A), consistent with recent findings^30^. Nevertheless, the number of cells undergoing programmed cell death was relatively low, suggesting that other mechanisms contribute more to the reduction in cell confluency. Therefore, we next decided to study the effect of RPB1 dosage on cell cycle progression, by pulsing cells with 5-ethynyl-2‘-deoxyuridine (EdU), and staining with an antibody against Cyclin E, followed by flow cytometry. Interestingly, even small reductions in RPB1 levels resulted in reduced incorporation of EdU (Figure 1I-J; Supplementary Figure S3C). Already when RPB1 levels were reduced by 30%, the fraction of cells in S-phase decreased measurably, with a concomitant increase in the fraction of cells in G1-phase (Figure 1I-J; Supplementary Figure S3B-C). Together, these data suggest that there is a critical threshold level of RPB1 required for the normal transition of cells from G1 to S-phase. Moreover, the surprisingly low depletion threshold again indicates that normal cellular functions are remarkably sensitive to RPB1 dosage, suggesting that RNAPII levels may be a crucial, limiting factor in determining transcriptional output.

### RPB1 levels are limiting for transcription of specific subsets of genes

To gain more insight into the importance of RPB1 dosage on nascent transcription, we next performed TT_chem_-seq^31,32^, using yeast 4TU-labelled RNA as a spike-in to control for the expected global changes in transcription. Remarkably, metagene plots indicated that even as little as 2.5% reduction in RPB1 levels results in reduced transcription levels genome-wide (Figure 2A). Consistent with our observation that 30-40% depletion of RPB1 (at 32-62 nM dTAG^V^-1) represents a critical threshold for cell proliferation, we found that this modest degree of RPB1 depletion indeed drastically reduces transcription (Figure 2A-C). Interestingly, stratification of all genes by biotype revealed that some gene classes are much more sensitive to RPB1 protein dosage than others. Most strikingly, a depletion of just 5% of RPB1 drastically reduced the transcription of small nuclear RNAs, while transcription of protein-coding genes and lncRNAs required somewhat higher levels of depletion for similar effects (Figure 2B; Supplementary Figure S4A).

**Figure 2.**
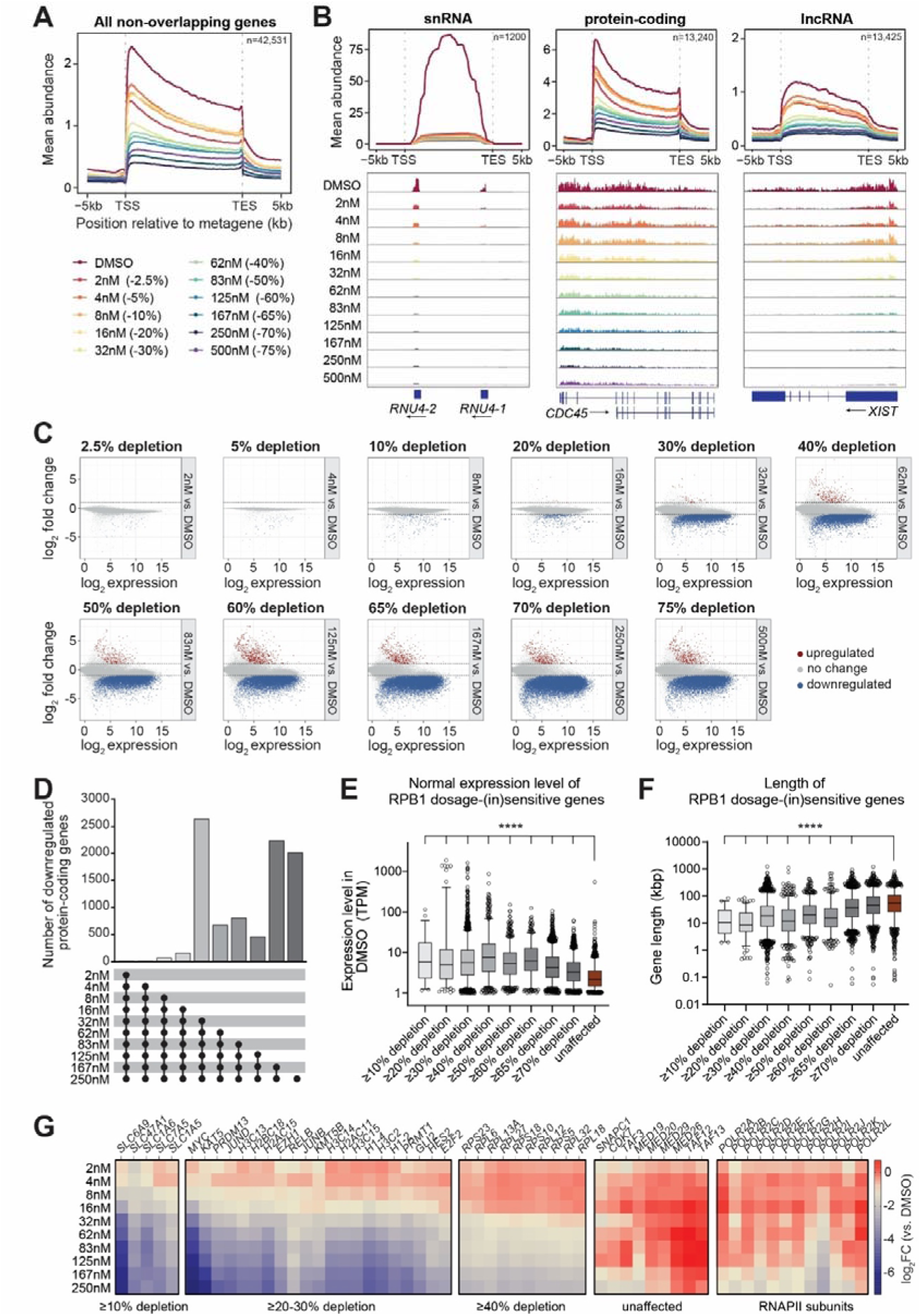
RPB1 levels are limiting for transcription of specific subsets of genes. **(A)** Metagene TT_chem_-seq profiles of all non-overlapping genes following treatment with different concentrations of dTAG^V^-1. Corresponding RPB1 depletion level is shown between brackets. Data are normalized to yeast spike-in. TSS=transcript start site. TES=transcript end site. Data represents mean of 3 replicates. **(B)** Metagene TT_chem_-seq profiles of non-overlapping genes stratified by biotype following treatment with different concentrations of dTAG^V^-1. Corresponding RPB1 depletion level is shown between brackets. Data are normalized to yeast spike-in. **(C)** Scatter plots showing the differentially expressed genes in the TT_chem_-seq data for all dTAG^V^-1 concentration vs. DMSO pairwise comparisons. Corresponding RPB1 depletion level is shown above in bold. Expression is basemean across all conditions, log2 fold-change has been shrunken using the apeglm method. Data are normalized to yeast spike-in. **(D)** Upset plot showing the number of protein-coding genes expressed above 1TPM that are consistently downregulated (hierarchical downregulation) in each dTAG^V^-1 concentration vs. DMSO pairwise comparison. Color-coding of bars is the same as used in (E) and (F). **(E)** Boxplots with baseline expression level (TPM value in DMSO condition) of genes found in each of the RPB1-dosage sensitivity bins. The genes in each bin are the same as in (D). Whiskers represent 5^th^-95^th^ percentiles. Unaffected genes are not differentially expressed in any of the dTAG^V^-1 vs. DMSO pairwise comparisons. Significance between unaffected genes and genes in the other RPB1 dosage-sensitivity bins was determined using a Kruskal-Wallis test. **(F)** Boxplots with length of genes found in each of the RPB1-dosage sensitivity bins. The genes in each bin are the same as in (D). Whiskers represent 5^th^-95^th^ percentiles. Unaffected genes are not differentially expressed in any of the dTAG^V^-1 vs. DMSO pairwise comparisons. Significance between unaffected genes and genes in the other RPB1 dosage-sensitivity bins was determined using a Kruskal-Wallis test. **(G)** Heatmap with log2 fold-change between each dTAG^V^-1 concentration vs. DMSO pairwise comparison for select genes from different RPB1 dosage-(in)sensitivity bin.

Further analysis of the TT_chem_-seq data revealed drastic rearrangement of the nascent transcriptome, with increasing dTAG^V^-1 concentrations resulting in more downregulated transcripts (Figure 2C). Even as little as a 20% reduction in total RPB1 levels resulted in a 2-fold downregulation of hundreds of genes, whereas a reduction of 30% resulted in downregulation of thousands of genes (Figure 2C; Supplementary Figure S4A; Supplementary Figure S4C). While at first it appeared that a small subset of genes was transcriptionally <u>up</u>regulated in response to RPB1 depletion (Figure 2C), manual inspection revealed that most of these were actually the consequence of transcription readthrough rather than increased transcription initiation (Supplementary Figure S4B), suggesting that, surprisingly, reduced RNAPII levels somehow decreases the efficiency of transcriptional termination.

Importantly, protein-coding genes that were downregulated upon mild RNAPII depletion conditions were also transcriptionally downregulated at higher levels of RNAPII loss (Figure 2D; Supplementary Figure S4C-F; see also Figure 2G). This indicates that the transcription changes follow a systematic hierarchy, with *additional* genes affected as RPB1 levels decrease, rather than being the consequence of stochastic changes, supporting the idea that transcription of numerous genes has evolved to be under fine control by RPB1 levels.

To better understand why the transcription of some genes was more sensitive to RPB1 dosage than others, we analyzed different features of protein-coding genes that were consistently affected at each of the RPB1 depletion levels (Figure 2D) and compared this to the subset of genes whose transcription was not significantly reduced upon depletion of RNAPII. This showed that the genes whose transcription was relatively unaffected even at the most extreme depletion levels were generally lowly expressed (Figure 2E). Moreover, we found that transcriptionally unaffected genes were also generally longer than RPB1 dosage-sensitive genes (Figure 2F). To gain more insight into the possible mechanisms behind the differential sensitivity of gene expression to RNAPII levels, we conducted a differential motif search in the core promoter of genes significantly downregulated by 30% RPB1 depletion, using the promoters of RPB1 dosage-<u>in</u>sensitive genes as background. Intriguingly, this approach yielded a consensus sequence of VRCCACGTGG as the most enriched motif, which closely resembles the binding motif of transcription factors n-MYC/c-MYC (Supplementary Figure S4G). Although it seems unlikely that loss of c-MYC alone can explain the reduced expression of all genes affected by 30% of RPB1 depletion, we note that the expression of *MYC* was indeed remarkably sensitive to RPB1 levels (Figure 2G and Supplementary Figure S4H).

Finally, we also found intriguing connections between dosage sensitivity and biological function. Specifically, a 20-30% depletion of RPB1 was sufficient to reduce the transcription of genes encoding histones, chromatin remodelers, and gene-specific transcription factors such as MYC (Figure 2G; Supplementary Figure S5). By contrast, genes encoding ribosomal proteins were generally unaffected at this point and instead required at least 40% RPB1 depletion before their transcription was altered (Figure 2G). Finally, we found it particularly interesting that the genes least affected by RPB1 depletion were enriched for genes associated with transcription initiation regulation and included subunits of the Mediator complex and general transcription factor TFIID (Figure 2G). Interestingly in this connection, transcription of several of the genes encoding RNAPII subunits was also well maintained, even when 75% of RNAPII in the cell was depleted (Figure 2G).

Together, these data point to an unexpected feedback mechanism by which cells preferentially protect expression of certain groups of genes when RNA polymerase II levels decrease. They also reveal that, on the other hand, genes known to be associated with rapid cell proliferation, such as histones and MYC, are remarkably sensitive to even slight changes in RNAPII levels.

### RPB1 concentration is a regulatory node for the recruitment of RNAPII to genes

The reduction in transcriptional activity observed by TT_chem_-seq might be the consequence of defects in any of the phases of the transcription cycle (Figure 3A). However, ultimately it is the recruitment of free RNAPII from the nucleoplasm to promoters in chromatin that initiates transcription. Indeed, the fact that highly expressed, short genes are more RPB1 dosage-sensitive than lowly transcribed, long genes would fit the idea that maintenance of the RNAPII pool is especially important for initiation (Figure 2E-F). To gain insight into the loading of RNAPII onto chromatin with changes in RPB1 dosage, we performed CUT&RUN^33^ using an anti-HA antibody to capture all forms of RNAPII, including the hypo-phosphorylated, pre-initiation form. We detected a clear RNAPII peak around the TSS of genes, which was gradually decreased upon increasing depletion of RPB1 (Figure 3B; Supplementary Figure S6A). This again suggests that the free RNAPII pool in the nucleoplasm is only just large enough to support normal transcription levels; as soon as RNAPII levels start decreasing, so does recruitment to promoters. Furthermore, this result opens the intriguing possibility that cellular transcription characteristics are not only determined by gene-specific transcription factors recruiting the general transcription machinery to promoters, but also by relatively modest changes in the abundance of RNAPII. In further agreement with this, we found that the population of initiating RNAPII molecules, marked by CTD serine 5 phosphorylation, was incrementally decreased with increasing dTAG^V^-1 doses (Figure 3C).

**Figure 3.**
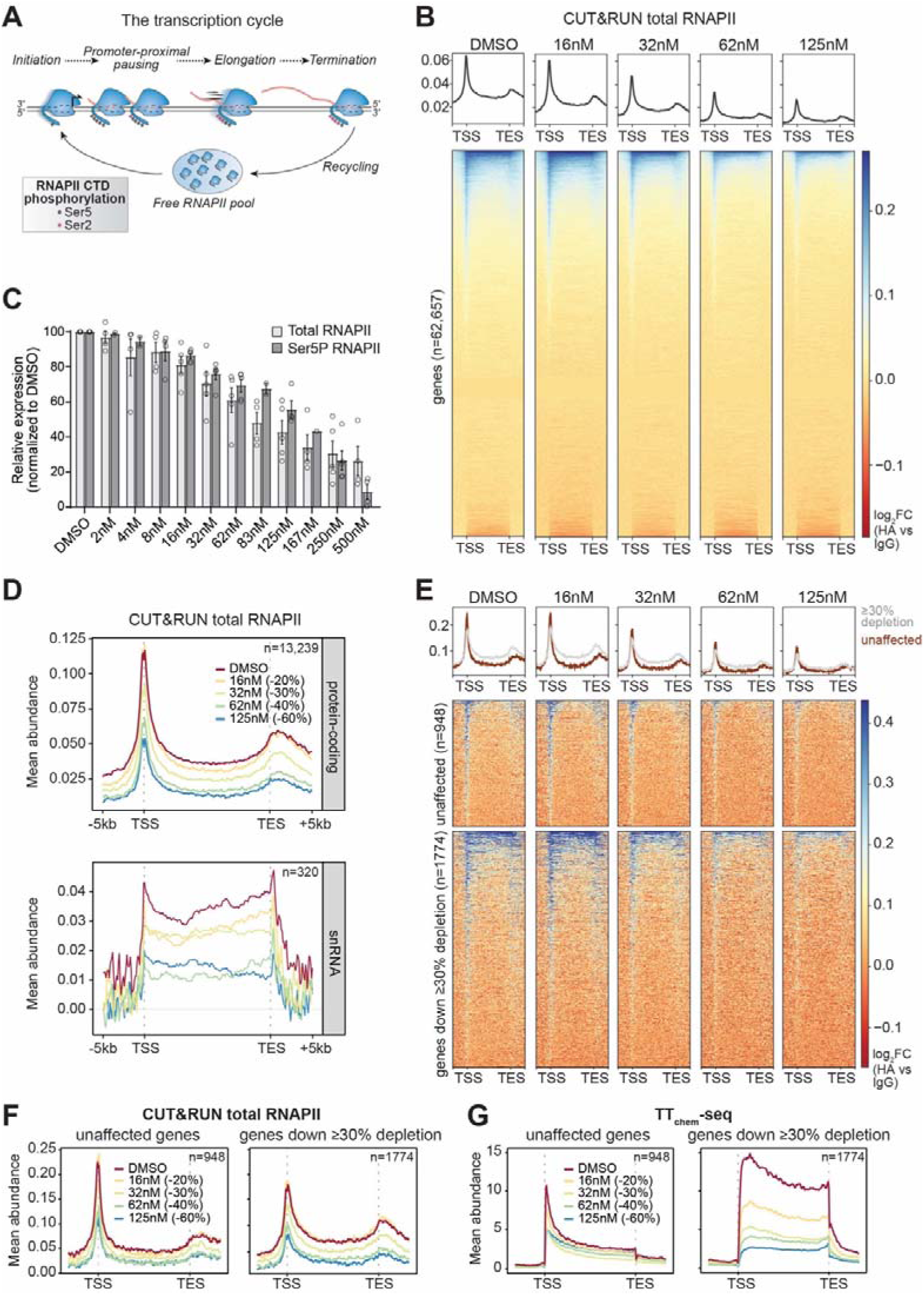
RPB1 concentration is a regulatory node for the recruitment of RNAPII to genes. **(A)** Schematic of the transcription cycle and phosphorylation status of the CTD of RPB1. **(B)** Metagene profiles (top) and heatmaps (bottom) of CUT&RUN data for total RNAPII using an antibody against HA-tag. Signal for RNAPII across all annotated genes is normalized to signal from IgG. TSS=transcript start site. TES=transcript end site. **(C)** Quantification of flow cytometry analysis of serine 5 phosphorylation on CTD of RPB1. Bargraphs represent mean of independent experiments ±SEM. **(D)** Metagene RNAPII CUT&RUN profiles of non-overlapping genes stratified by biotype following treatment with different concentrations of dTAG^V^-1. Corresponding RPB1 depletion level is shown between brackets. Signal for total RNAPII across genes is normalized to signal from IgG. **(E)** Metagene profiles (top) and heatmaps (bottom) of CUT&RUN data for total RNAPII (normalized to IgG). RNAPII signal across unaffected genes and genes consistently downregulated with ≥30% depletion of RPB1 are shown at the different dTAG^V^-1 concentration treatments. Genes in these bins are the same as Figure 2D, except that overlapping genes have been removed. **(F)** Metagene RNAPII CUT&RUN profiles for unaffected genes and genes consistently downregulated with ≥30% depletion of RPB1 following treatment with different concentrations of dTAG^V^-1. Corresponding RPB1 depletion level is shown between brackets. **(G)** Metagene TT_chem_-seq profiles for unaffected genes and genes consistently downregulated with ≥30% depletion of RPB1 following treatment with different concentrations of dTAG^V^-1. Corresponding RPB1 depletion level is shown between brackets.

To determine whether the loading of RNAPII onto chromatin is affected uniformly or whether reduced transcription initiation contributes to the differential sensitivity of genes to RPB1 dosage, we next stratified genes by gene type. Consistent with the strong sensitivity of snRNA transcription to RNAPII levels, we found that relatively small depletions of RPB1 affected the loading of RNAPII onto snRNA promoters much more than at protein-coding genes (Figure 3D). This supports the idea that RPB1 dosage can act as a regulatory node for the recruitment of the polymerase to specific genes.

In extension of this, we sought to determine whether the differential sensitivity of protein-coding genes to RPB1 dosage might be attributed simply to reduced transcription initiation. Intriguingly, incremental depletion of RPB1 resulted in a gradual reduction of RNAPII occupancy both on the genes that were significantly downregulated by 30% depletion and on the RPB1 dosage-<u>in</u>sensitive genes (Figure 3E-G). The fact that the loading of RNAPII is affected in both gene classes suggests that other mechanisms in addition to initiation must be at play to explain their differential sensitivity to RPB1 dosage. Indeed, somewhat surprisingly, RNAPII occupancy around the TSS was generally slightly higher for RPB1 dosage-<u>in</u>sensitive genes than at the genes that were markedly downregulated upon 30% RNAPII depletion (Figure 3E), despite the generally lower transcription levels of the former genes (Figure 2E; Figure 3G). This again points to post-initiation effects, such as at promoter-proximal pausing.

### Reduced promoter-proximal pausing compensates for RNAPII depletion

Upon initiation, RNAPII typically pauses ∼60 bp into a gene, at promotor-proximal pause sites, until the full elongation complex has been correctly assembled and the nascent RNA has been capped^16^. To investigate promoter-proximal pausing, we performed precision run-on sequencing (PRO-seq^34^) which maps the location of paused polymerases with nucleotide resolution. As expected, metagene plots from these experiments revealed a prominent peak at the 5’ end of genes, corresponding to the location of polymerases at the promoter-proximal pause (Figure 4A). Intriguingly, we found that this peak was drastically reduced with decreasing RPB1 levels in the cell, while the coverage in the gene body was relatively unaffected. We therefore calculated the pausing index, which provides a measure of promoter-proximal pausing^34^. This revealed that incremental decreases in RPB1 levels lead to lower pausing indices, suggesting that RNAPII pauses less, *i.e.* is released more efficiently into the gene body when the RNAPII pool is low (Figure 4B). This points to the existence of a feedback loop whereby reduced recruitment of RNAPII to transcription initiation sites is balanced by changes in promoter-proximal pausing or reduced premature termination. Release of RNAPII from promoter-proximal pause sites is accompanied by increased CTD phosphorylation, at both serine 5 and serine 2. Indeed, using flow cytometry, we found that the population of Ser5+2 phosphorylated RNAPII (marking elongating RNAPII^35^) was indeed less reduced than expected from the reduction in total RNAPII (Supplementary Figure S7A). Moreover, CUT&RUN revealed an *increase* in the genome-wide occupancy of Ser5+2P RNAPII on protein-coding genes in response to RPB1 depletion, even though the overall occupancy of RNAPII was decreased (Figure 4C; Figure 3B). In other words, the fraction of RNAPII engaged in elongation increases upon RPB1 depletion. Together, these data support a model in which reduced RPB1 levels result in rapid release from the promoter-proximal pause to counteract the lower RNAPII recruitment to promoters. Importantly, RNAPII is released more efficiently from the promoter-proximal pause region of both RPB1 dosage-sensitive and-insensitive genes (Supplemental Figure S7B). However, the pausing index of RPB1 dosage-insensitive genes is generally much higher than that of RPB1 dosage-sensitive genes (Supplemental Figure S7C), suggesting that promoter-proximal pausing might normally be an important regulatory checkpoint for RPB1 dosage-insensitive genes.

**Figure 4.**
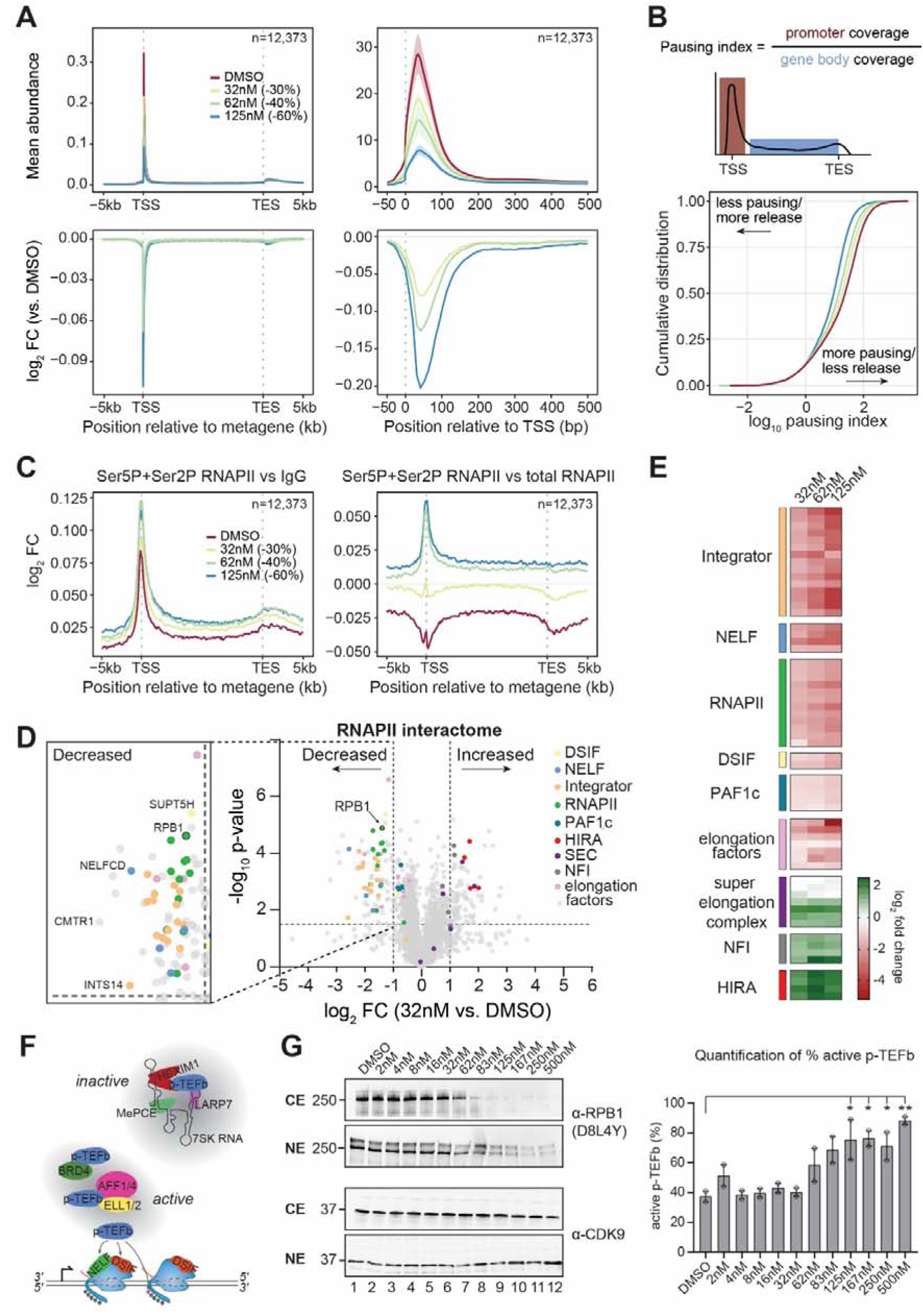
Reduced promoter-proximal pausing helps maintain transcriptional activity upon RNAPII depletion. **(A)** PRO-seq profiles (top) and log2 fold-change profiles (dTAG^V^-1 vs. DMSO) (bottom) of all non-overlapping protein-coding genes >2kb following treatment with different concentrations of dTAG^V^-1. Corresponding RPB1 depletion level is shown between brackets. Metagene profiles are shown on the left, profiles aligned at the TSS are shown on the right. Data are normalized to drosophila spike-in. Data represents mean of 3 replicates ±SEM. **(B)** Schematic showing pausing index calculation (top) and empirical cumulative distribution function (ECDF) plot of pausing index of all non-overlapping protein-coding genes >2kb following treatment with different concentrations of dTAG^V^-1. **(C)** Metagene CUT&RUN profiles for serine 5 + serine 2 phosphorylated (Ser5P+Ser2P) RNAPII across all non-overlapping protein-coding genes following treatment with different concentrations of dTAG^V^-1. Corresponding RPB1 depletion level is shown between brackets. Signal for Ser5P+Ser2P RNAPII is either normalized to signal from IgG (left) or total RNAPII (right). **(D)** The change in RNAPII interactome following 30% depletion of RPB1. Factors enriched following RPB1 depletion are shown on the right, while factors that are depleted from RNAPII following RPB1 depletion are shown on the left (enlarged and labeled on the left). **(E)** Heatmap of known RNAPII interactors with log2 fold changes following treatment with different dTAG^V^-1 concentrations compared to DMSO. **(F)** Schematic of different subcomplexes containing pTEFb. **(G)** Western blot analysis of active (NE) and inactive (CE) CDK9 following treatment with different dTAG^V^-1 concentrations. Blot for total RPB1 serves as control for effective dTAGV-1 treatment and fractionation. Quantification of the percentage active pTEFb from two independent experiments is shown on the right. Data is represented as mean ±SD. Statistical significance between DMSO and dTAG^V^-1 conditions was determined using a mixed-effects model, followed by Dunnett’s multiple comparisons test. *=P-value<0.05; **=P-value<0.01.

In an attempt to gain further insight into the mechanism underlying this transcriptional feedback loop, we determined the RNAPII interactome by subjecting immunoprecipitates of hyper-phosphorylated, transcriptionally engaged RNAPII to mass spectrometry. As expected, the addition of dTAG^V-^1 reduced the amount of RNAPII immunoprecipitated, resulting in lower detection of RNAPII subunits in these conditions (Figure 4D; Supplementary Figure S7D). Remarkably, however, components of the pausing-specific NELF and Integrator complexes were reduced even more drastically than RPB1 after addition of 32nM, 62nM and 125nM dTAG^V^-1 (Figure 4D-E; Supplementary Figure S7D). In other words, RNAPII complexes containing NELF and integrator were reduced relative to total RNAPII. Given its role in dissociating paused RNAPII, the reduced abundance of Integrator-bound RNAPII complexes after dTAG^V^-1 treatment is compatible with a model in which depletion of RNAPII leads to less premature termination at the promoter-proximal pause. Similarly, the reduced number of NELF-bound RNAPII complexes is consistent with more rapid release from promoter-proximal pausing suggested by PRO-seq and CUT&RUN for Ser5+Ser2 phosphorylated RNAPII (Figure 4A-E).

Conversely, we noticed that depletion of RNAPII resulted in markedly increased association with the p-TEFb kinase, a crucial, positive regulator of promoter proximal pausing (Figure 4D-E; Supplementary Figure S7D-S6E). The activity of p-TEFb is governed by its reversible association with different binding partners: P-TEFb is part of a ‘super elongation complex’, and bound by transcription factors such as BRD4, while sequestration by the 7SK snRNP renders it inactive in a high molecular weight complex (see Figure 4F)^36^. Interestingly, the IP-MS data also revealed an increased interaction between RNAPII and other subunits of the super elongation complex (AFF1/AFF4, ELL/ELL2) upon dTAG^V^-1 treatment, supporting the idea that RPB1 depletion increases p-TEFb activity. Indeed, separation of the large (inactive) and free (active) p-TEFb forms using differential salt extractions showed that increasing concentrations of dTAG^V^-1 resulted in a reduction of soluble (inactive) pTEFb, while the amount of active p-TEFb increased (Figure 4G). Thus, reduced RPB1 levels lead to increased pTEFb activity, somewhat akin to how inhibition of transcription by pTEFb inhibition or DNA damage promotes the release of pTEFb from 7SK snRNP^37–39^.

### Reduced RPB1 levels results in altered transcript elongation rates

Other factors that were co-immunoprecipitated with RNAPII at significantly higher levels when RNAPII dosage was decreased included components of the HIRA complex (HIRA, UBN1/UBN2, CABIN1) (Figure 4D-E, Supplementary Figure S7D). The HIRA histone chaperone complex incorporates histone variant H3.3 in euchromatin in a replication-independent manner, and H3.3 levels have previously been correlated with RNAPII elongation rates^40–43^. An increased association of RNAPII with HIRA might therefore point to an effect of RNAPII depletion on elongation rates as well. As an initial indication of such changes, difference maps revealed that increasing dTAG^V^-1 concentrations generally affected transcription activity at the end of genes less than would be expected from the decreased transcriptional activity observed at the beginning of genes (Figure 5A). Indeed, further analysis of the TT_chem_-seq data revealed an increased elongation index in long genes when RPB1 abundance is low (Figure 5B).

**Figure 5.**
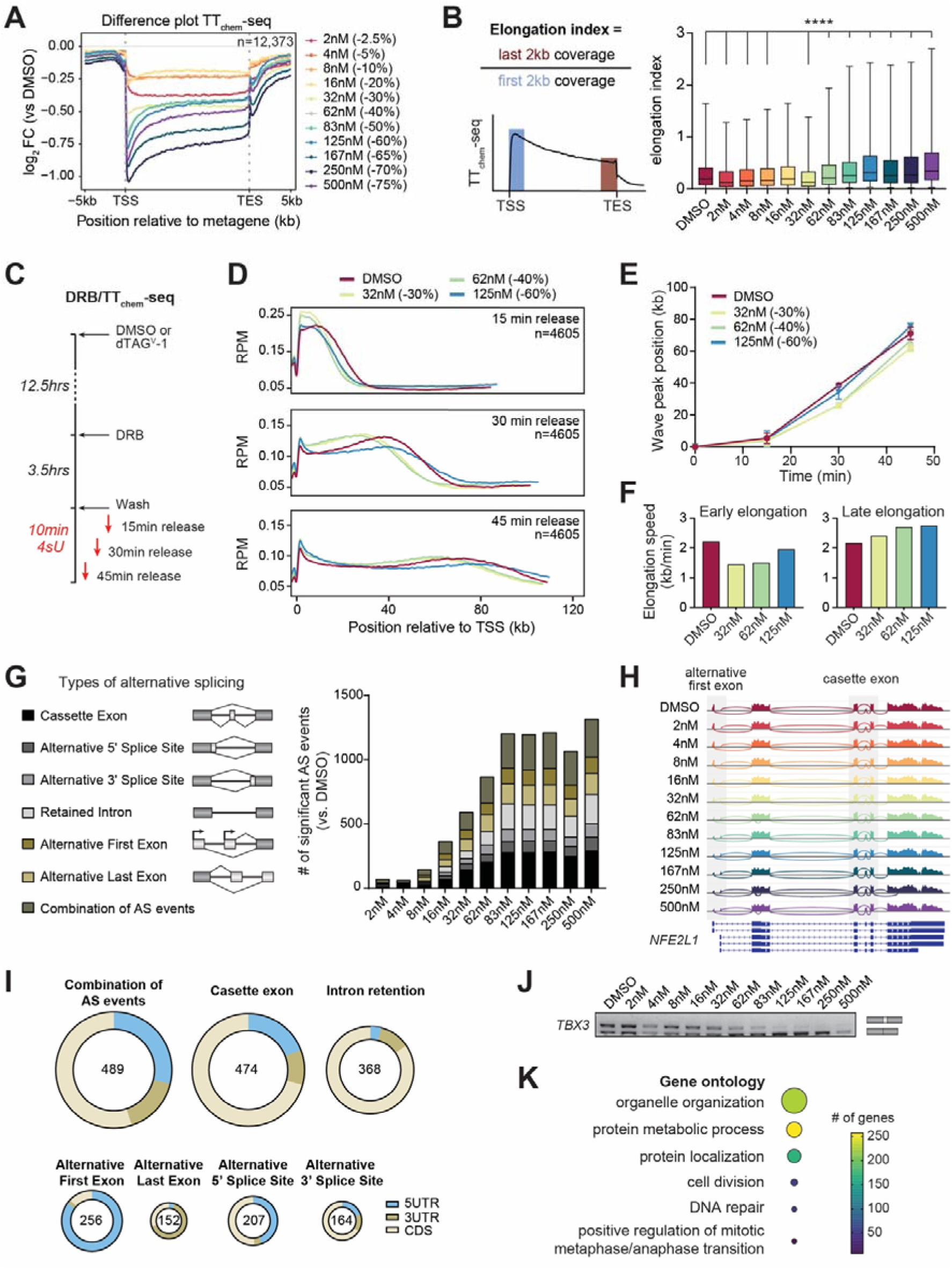
Reduced RPB1 levels results in altered transcript elongation rates and splicing. **(A)** Metagene TT_chem_-seq profiles of all non-overlapping protein-coding genes with log2 fold change in signal following treatment with different concentrations of dTAG^V^-1 compared to DMSO. Corresponding RPB1 depletion level is shown between brackets. Data are normalized to yeast spike-in. TSS=transcript start site. TES=transcript end site. Data represents mean of 3 replicates. **(B)** Schematic showing elongation index calculation (left) and boxplots of elongation index of all non-overlapping protein-coding genes >2kb following treatment with different concentrations of dTAG^V^-1. Whiskers represent 5^th^-95^th^ percentiles. Statistical significance between DMSO and dTAGV-1 treatment conditions was determined using a Friedman test, followed by Dunn’s multiple comparisons test. ****=P-value<0.0001. **(C)** Schematic of the DRB/ TT_chem_-seq workflow. **(D)** Metagene plots of RPM-normalized TT_chem_-seq profiles for non-overlapping protein-coding genes ≥60kb and ≤300kb following 15, 30 or 45 minutes release from DRB. (E) Location of the wave peak following different times of DRB release following treatment with different concentrations of dTAG^V^-1. Corresponding RPB1 depletion level is shown between brackets. **(F)** Bargraphs with average elongation speed following treatment with different concentrations of dTAG^V^-1 (3 replicates) across all non-overlapping protein-coding genes ≥60kb and ≤300kb. Early elongation speed is determined between 15 and 30 minutes release from DRB, whereas late elongation speed is calculated from the distance traveled between 30 and 45 minutes release from DRB. **(G)** Schematic of the different types of alternative splicing events (left) detected following treatment with different concentrations of dTAG^V^-1 (right). Significant AS events were defined as |PSI-value|≥0.1 and P-value<0.05. **(H)** Sashimi plot for NFE2L1, showing two representative examples of alternative splicing (alternative first exon and cassette exon), following treatment with different concentrations of dTAG^V^-1. **(I)** Doughnut plots with distribution of AS events affecting the 5’ untranslated region (UTR), coding sequence (CDS) and 3’UTR of protein-coding genes. Total number of AS events in each category shown in the middle. **(J)** RT-PCR showing increased skipping of exon 2a in *TBX3* upon dTAG^V^-1 treatment, which is also alternatively spliced in response to altered elongation rates. **(K)** Gene ontology driver terms enriched for by alternatively spliced protein-coding genes.

To directly test whether transcript elongation is influenced by RNAPII dosage, we next used DRB/TT_chem_-seq^31,32^. In this approach, DRB is used to inhibit the kinase activity of CDK9 in P-TEFb, preventing escape from promoter-proximal pausing and restricting RNAPII transcription to the first few hundred bases of the gene, while allowing already transcribing polymerases to run off the genes. Upon subsequent DRB removal, RNAPII is released and resumes transcript elongation across genes across the genome (Figure 5C). Interestingly, 30-40% depletion of RPB1 resulted in a reduction in the progress of elongating RNAPII at all timepoints (Figure 5D). Upon ∼60% depletion of total RNAPII, slower elongation was observed in the first 15 minutes after DRB release, but at later times slightly faster elongation was observed (Figure 5E). These intriguing effects on elongation were slight but reproducible. Calculation of the elongation rates using the position of the wave peak across time showed that the speed of RNAPII was lower close to the promoter after RPB1 depletion but higher than normal further into the gene body (Figure 5F). These data thus suggest the presence of another feedback loop in which reductions in RNAPII abundance trigger changes in the nature of elongation of the remaining polymerases.

### Splicing changes upon RNAPII depletion

The elongation rate of RNA polymerase II influences several co-transcriptional processes, such as splicing, polyadenylation, and transcription termination^44^. Slowing down of RNA polymerases at the end of genes is important for proper transcription termination, which is why stimuli that increase elongation speed frequently result in transcription read-through. Indeed, we note that the increased elongation speed in response to 60% depletion in RPB1 was correlated with increased levels of transcription read-through (*cf.* Supplementary Figure S4B). Given that changes in transcript elongation affect splicing and that the transcription levels of spliceosomal snRNAs were particularly sensitive to RPB1 dosage (*cf.* Figure 2B), we also investigated the effects of RPB1 dosage on mRNA splicing. To this end, we performed mRNA-seq and analyzed changes in mRNA splicing as well as in the usage of alternative polyadenylation sites. These experiments showed that increased depletion of RPB1 results in gradually increased levels of alternative splicing (Figure 5G). Depletion of RNAPII to around 50% of normal levels resulted in more than a thousand splicing changes, but further depletion did not result in additional changes, suggesting that, intriguingly, splicing fidelity is primarily affected within a defined range of relatively small changes in RPB1 dosage and transcriptional output. We note that while the reduction in snRNAs upon RNAPII depletion might be expected to result in a general decrease in splicing capability and thus in the retention of introns, widespread evidence of such an effect was not observed (Figure 5G). Instead, we predominately detected increased usage of cassette exons upon RPB1 depletion, giving rise to alternative mRNA isoforms (Figure 5G-J). Finally, we found that moderate reductions in RPB1 levels led to increased usage of alternative first and last exons (Figure 5G-H). Intriguingly, many of these events affected the 5’UTR and 3’UTR rather than the coding sequence of genes (Figure 5I), opening the possibility that RPB1 dosage modulates mRNA stability and/or translation efficiency of these genes at least partly through changes in alternative splicing.

### Increased mRNA stability in response to reduced transcription levels

The results above indicate that changes in RNAPII dosage have a marked effect on basic transcription sub-processes, mRNA splicing, and even termination, and indicate the existence of cellular buffering or feedback loops to mitigate these changes. To determine whether and how RNAPII dosage affects the steady-state transcriptome, we next analyzed the mRNA-seq data for differential gene expression. Surprisingly, depletion of RNAPII not only resulted in reduced levels of many mRNAs, but also upregulation of hundreds of mRNA transcripts (Figure 6A). Importantly, while transcription was indeed down-regulated at the vast majority of genes that encode downregulated mRNA transcripts, as expected, most of the upregulated mRNAs were encoded by genes whose transcription was not markedly affected by reduced RPB1 levels (Figure 6B). The most obvious explanation for this discrepancy is that reduced RNAPII dosage results in decreased decay of many mRNAs. In apparent agreement with this interpretation, we did not observe a significant overall change in the total RNA mass after RPB1 depletion (Figure 6C) and indeed observed an increased signal when using oligo-dT probes combined with fluorescent in situ hybridization to measure total mRNA level (Figure 6D). To investigate further, we calculated the half-life of mRNA by determining the ratio of expression measured by mRNA-seq relative to the TT_chem_-seq levels, for all protein-coding genes. mRNA half-life was anti-correlated with RNAPII levels, peaking in a two-fold average increase in mRNA half-lives at very high levels of RNAPII depletion (70%; 250nM dTAG^V^-1) (Figure 6E). Thus, our data suggests that mRNA stability is increased in response to reduced RPB1 levels, resulting in buffering of mRNA concentration.

**Figure 6.**
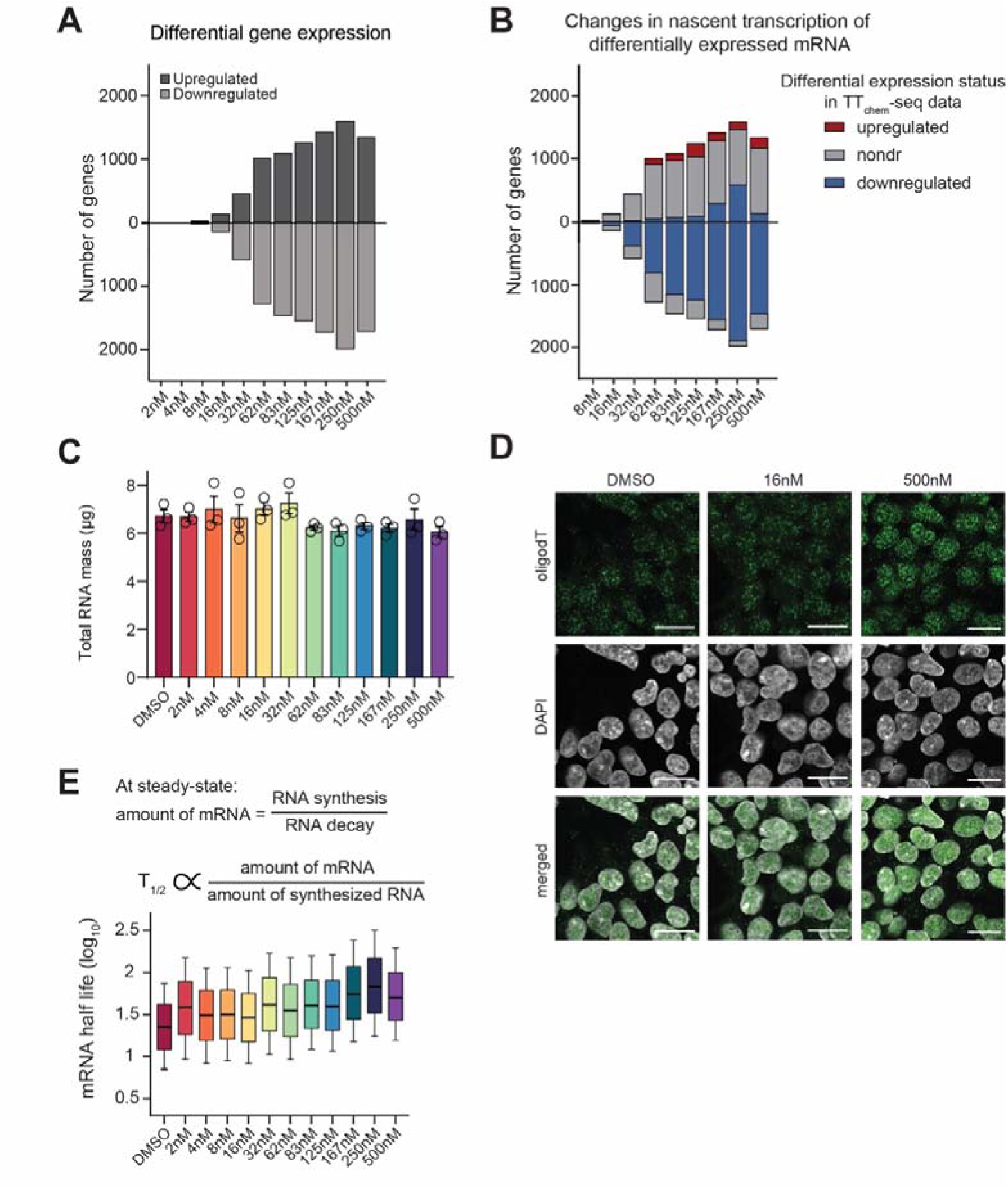
Increased mRNA stability in response to reduced transcription levels. **(A)** Bar graphs of genes differentially expressed at steady-state following treatment with different concentrations of dTAG^V^-1 compared to DMSO. Positive side of y-axis: upregulated genes in mRNA-seq data, negative side of y-axis: downregulated genes in mRNA-seq data. **(B)** Bar graphs showing the differential expression status by TTchem-seq analysis for genes differentially expressed at steady-state following treatment with different concentrations of dTAG^V^-1 compared to DMSO. Positive side of y-axis: upregulated genes in mRNA-seq data, negative side of y-axis: downregulated genes in mRNA-seq data. **(C)** Quantification of total RNA mass extracted from cells following treatment with different concentrations of dTAG^V^-1. Data represents mean of 3 replicates ±SEM. **(D)** Fluorescent in situ hybridization with an oligodT probe showing poly(A)+ mRNA following treatment with 16nM and 500nM dTAG^V^-1. Nuclei are marked with DAPI. Scale bar=20um. **(E)** Calculation of mRNA half-life (top) following treatment with different concentrations of dTAG^V^-1. Data is shown as boxplots, with whiskers representing 5^th^-95^th^ percentiles.

The results shown so far support the conclusion that the cellular levels of RNAPII are only just high enough to maintain transcription. Conversely, increases in RPB1 levels might also affect transcription. To test this possibility, we expressed an RPB1 transgene under the strong, constitutive *UBC* promoter in the RPB1 degron cell line. Stable expression of this transgene was enough to restore RPB1 protein levels to normal levels during treatment with 500 nM dTAG^V^-1 (Supplementary Figure S8A-B). Intriguingly, however, in DMSO conditions where endogenously expressed RPB1 was not degraded, expression of the RPB1 transgene did not result in the expected doubling of RPB1 protein levels but only resulted in slight (∼30%) overexpression (Supplementary Figure S8A-B). Nevertheless, this relatively small overexpression resulted in a modest increase in nascent transcription genome-wide (Figure 7A). Moreover, it resulted in a 2-fold increase in the transcription of snRNAs and histones, the genes that were also most sensitive to depletion of RPB1 (Supplementary Figure S8C). At the steady-state level, we found that expression from the transgene alone rescued the effects of RPB1 dTAG depletion on mRNA half-life (Supplemental Figure S8D, compare the 500nM graphs), while the increase in RPB1 levels in DMSO conditions resulted in an overall decrease in mRNA half-life (Supplemental Figure S8D).

**Figure 7.**
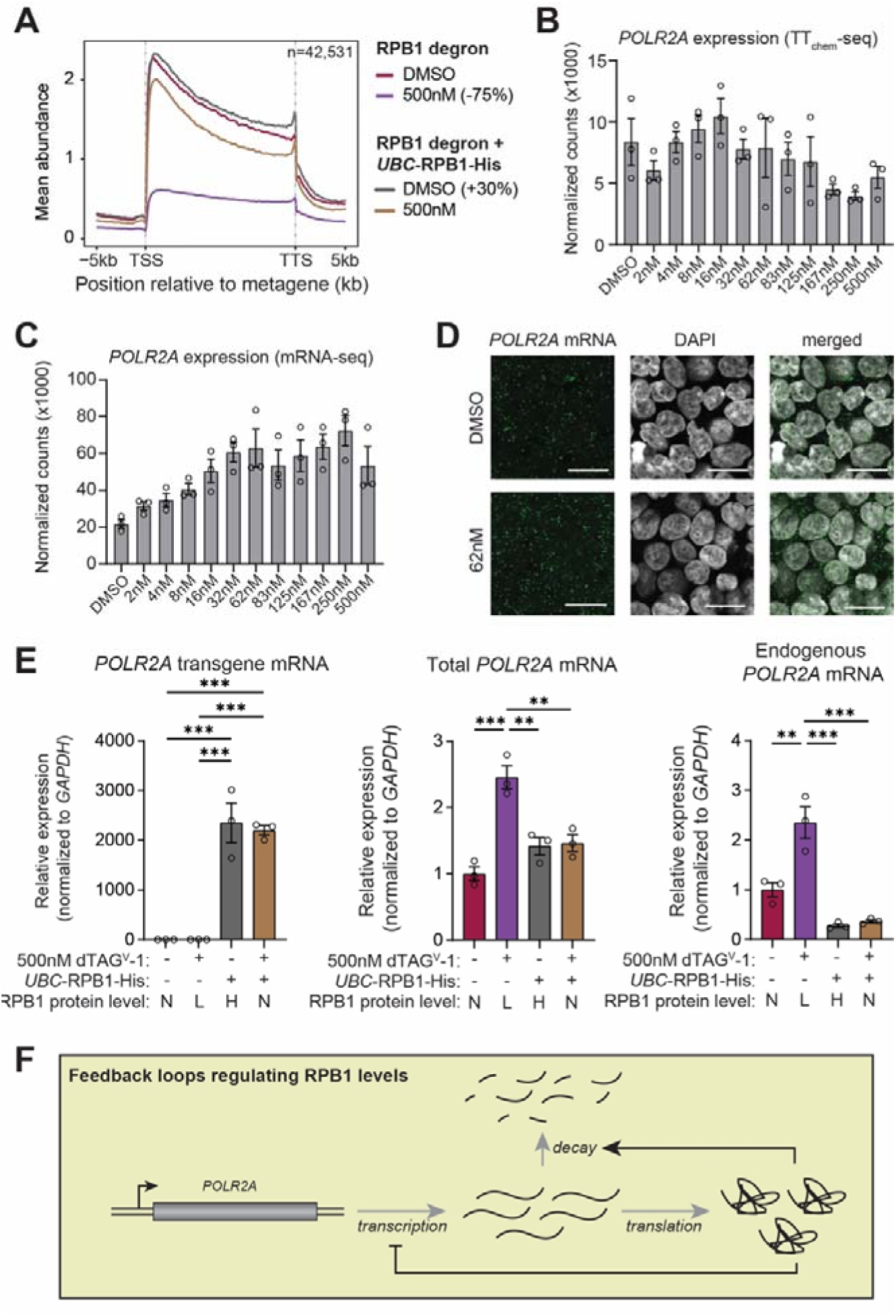
Regulation of *RPB1* expression by RPB1. **(A)** Metagene TT_chem_-seq profiles of all non-overlapping genes following expression of the RPB1 transgene from a *UBC*-promoter. Corresponding RPB1 levels in each condition is shown between brackets. Data are normalized to yeast spike-in. TSS=transcript start site. TES=transcript end site. Data represents mean of 3 replicates. **(B)** Bar graphs with expression of nascent *POLR2A* RNA as determined by TTchem-seq, following treatment with different concentrations of dTAG^V^-1. **(C)** Bar graphs with expression of *POLR2A* mRNA as determined by mRNA-seq, following treatment with different concentrations of dTAG^V^-1. **(D)** Fluorescent in situ hybridization for *POLR2A* mRNA following treatment with 62nM dTAG^V^-1. Nuclei are marked with DAPI. Scale bar=20um. **(E)** qRT-PCR results for the relative expression of *POLR2A* following 500nM dTAG^V^-1 treatment in the 6XG linker RPB1 degron cell line and the stable RPB1 degron cell line expressing an RPB1 transgene from a *UBC* promoter. Expression for *POLR2A* transgene (left), total *POLR2A* mRNA (middle) and *POLR2A* RNA transcribed from the endogenous locus (right) are normalized to *GAPDH*. Expression is set to 1 in RPB1 degron cells following DMSO treatment. The relative RPB1 protein levels are presented below: N=normal; L=low; H=high. Statistical significance was determined using one-way ANOVA, followed by post-hoc Tukey test. **=P-value<0.01; ***=P-value<0.005. **(F)** Schematic depicting the feedback loops from RPB1 protein on the synthesis and turnover of its own mRNA.

### Regulation of *RPB1* expression by RPB1

Together, the findings above indicate that RPB1 levels are limiting for transcription activity but also open the possibility that they are closely monitored and controlled by the cell to enable finely tuned gene expression. In support of this idea, we found that while nascent transcription of the *POLR2A* gene itself unsurprisingly decreases at very high levels of RPB1 loss, smaller, more physiologically relevant changes in RPB1 levels (up to 20% depletion; 16 nM dTAG^V^-1) resulted in somewhat <u>in</u>creased transcription of its own locus (Figure 7B), and, more importantly, that upon gradual depletion of the RPB1 protein (up to 40% depletion; 62 nM dTAG^V^-1), ∼three-fold more RPB1 mRNA was detected by both mRNA-seq (Figure 7C) and quantitative reverse transcriptase PCR (qRT-PCR) (Supplementary Figure S7E). Fluorescent *in situ* hybridization for *POLR2A* mRNA confirmed the increased expression without a noticeable change in its subcellular distribution (Figure 7D).

To further investigate the possible existence of a feedback loop whereby the RPB1 protein regulates both the synthesis and stability of its own mRNA, we now used the degron cell line that also expresses RPB1 from a *UBC* promoter-driven transgene, in combination with qRT-PCR and primers that can distinguish the *POLR2A* mRNA forms expressed from the endogenous promoter and the transgene promoter, respectively. As expected, the transgene-specific *POLR2A* mRNA was only detected in the cell line expressing the *UBC*-driven construct (Figure 7E, left). Moreover, the increase in endogenous RPB1 mRNA observed above by mRNA-Seq after RPB1 protein degradation was clearly visible by qRT-PCR of total RPB1 mRNA as well (Figure 7E, middle, 2 bars on left). More importantly, this increase was largely annulled by overexpression of RPB1 from the transgene, with total mRNA levels only moderately above normal levels even when both *POLR2A* and the *UBC*-driven transgene were present (Figure 7E, middle, 2 bars on right). Indeed, expression of mRNA encoding RPB1 specifically from the endogenous *POLR2A* locus was dramatically reduced in the cell line containing the *UBC*-driven transgene (Figure 7E, right).

Together, these results provide evidence that RPB1 protein levels regulate the level of (RPB1-expressing) *POLR2A* mRNA (Figure 7F).

## Discussion

Maintaining protein concentration within a homeostatic range helps protect cells against perturbations such as environmental changes or noise in biochemical reactions^45^. However, for most proteins, the permissible limits of this range remain unclear. Here, we investigated the sensitivity of human HEK293 cells to RNAPII (RPB1) dosage and found that only minimal reductions can be tolerated before cell growth and progress through the cell cycle is perturbed. Conversely, despite a 7-fold overexpression at the *RPB1* mRNA level, the RPB1 protein levels could not be <u>in</u>creased beyond 130% of baseline, reinforcing the notion that human cells only tolerate a narrow range of RPB1 protein abundance. Our data also indicate that RNAPII levels are normally maintained within an acceptable range by the RPB1 protein regulating *RPB1* (*POLR2A*) mRNA expression. Finally, we also established that when RPB1 abundance moves out of the acceptable range, this has dramatic effects on all subprocesses of the transcription cycle, as well as mRNA splicing and stability. Indeed, meaningful, dynamic feedback loops between the processes have clearly evolved.

### Why are RNAPII levels maintained in a narrow range?

An important question that arises from these findings is why cellular RPB1 levels are maintained in such a narrow range. The fact that even small decreases in RPB1 dosage have radical effects on transcription and cell proliferation thus makes it obvious to ask *why* this fundamental cellular characteristic evolved. One possibility is that producing excess protein has in general been selected against in evolution. In the case of a heteromultimeric protein complex such as RNAPII, where the abundance of individual subunits needs to be balanced to maintain proper stoichiometry, this evolutionary drive might be especially pertinent. However, we propose that constraints on RPB1 abundance also evolved because there is a significant cellular benefit to maintaining RPB1 at near-limiting levels: the narrow abundance range with even small changes in RPB1 levels resulting in broad and non-stochastic changes in transcriptional output may thus enable cells not only to maintain homeostasis but also to respond to intra-and extracellular cues and adjust to new conditions. Intrinsic in this working model is that specific genes evolved to respond, or not respond, to changes in RNAPII levels. They do so for a purpose, or at least non-randomly. This idea is supported by even a cursory glance at the genes in each category, with – for example - genes encoding factors needed to maintain transcriptional initiation being relative robustly expressed even at low levels of RNAPII, and those encoding factors promoting cellular growth such as MYC being extremely sensitive. Remarkably, an obvious prospect raised by these results is that the hallmarks of different cell types might be determined not only by the transcription factor make-up but also by cell-type specific RNAPII levels.

Interestingly, the idea that cells dynamically regulate RPB1 levels in response to cellular cues may also help to explain the puzzling finding that so many different E3 ubiquitin ligases, including Cullin Ring Ligase 3 with ARMC5 (CRL3^ARMC^^5^), CRL4^CSA^, CRL4^ElonginA^ and CRL2^VHL^ all target RPB1 for poly-ubiquitylation and degradation in a variety of distinct cellular response pathways^46–51^. One extreme example is UV-irradiation which results in transcription being rapidly shut down in a global manner. Ubiquitylation and degradation of RPB1 is critical for this shutdown^26^. Together with the data reported here, this supports a model in which RPB1 production and degradation are tightly interwoven to dynamically optimize cellular transcription homeostasis in the face of changes to the intra-or extracellular milieu.

Conversely, transcription output also needs to be globally <u>in</u>creased in response to increases in cell size, so as to maintain mRNA concentration. Cells frequently change in size, for instance while progressing through the cell cycle. Interestingly, it was recently shown in yeast that simultaneous overexpression of all RNAPII subunits resulted in increased loading of RNAPII on chromatin, whereas increased expression of general transcription factors did not^21^. This suggests that RNAPII abundance may be a rate-limiting component for transcription initiation also during cell size scaling in yeast. While overexpression of RPB1 may not *in itself* be sufficient to drive changes in cell size, as this may require growth factors and other signals, loss of RPB1 would be predicted to result in a breakdown of biosynthetic scaling. Such a breakdown has previously been shown to drive cells towards a permanent exit of the cell cycle, and entrance into senescence^52–56^. We found that even small reductions in human RPB1 levels result in an accumulation of cells in G1/G0. Moreover, others have shown that knockdown of *POLR2A* in human skin fibroblasts increases cellular senescence^57^ and certain mutations in the *RPB1/POLR2A* gene (*RPO21* in yeast) result in G1 arrest in both yeast and humans^58,59^. This supports the idea that cells need a certain minimal transcription capacity to progress through the G1 restriction point and that they therefore exit the cell cycle when RNAPII levels are too low. This also raises the exciting possibility that decreased RPB1 levels may act as a regulatory node to control cell cycle exit, such as during neurogenesis, myogenesis or erythropoiesis.

### Regulation of RNAPII levels

The narrow tolerable range of RPB1 abundance in human cells suggests that synthesis and decay of RPB1 (and in turn RNAPII) must be balanced precisely. Intriguingly, pervasive degradation of RPB1 under homeostatic cell growth conditions was recently shown to be mediated by the E3 ubiquitin ligase ARMC5^49,51,60,61^. Our data reveal that the synthesis of RPB1 is also heavily regulated. Intriguingly, we thus found that the levels of RPB1 mRNA are regulated through a negative feedback loop involving the RPB1 protein. This feedback loop, in which the RPB1 protein appears to both inhibit transcription of its own encoding gene and promote decay of its own mRNA, helps ensure that RPB1 levels are maintained within its homeostatic range. Inherent to such a feedback loop is the existence of a ‘sensor’ of RNAPII abundance. It has recently been proposed that the splicing factor PTBP1 communicates information about RNAPII levels from the nucleus to the cytoplasm to activate apoptotic signaling and might thus function as such a sensor^62^. Moreover, previous research indicated that nuclear mRNA abundance negatively regulates RNAPII abundance^25^. Understanding the mechanisms that underlie the intriguing feedback loops governing RPB1 abundance thus represents an important future goal.

### Feedback loops that buffer transcriptional output

Our data indicate that, not surprisingly, lower RNAPII levels also result in less loading of the polymerase onto promoters and thus presumably less transcription initiation. Interestingly, however, at many genes these effects appear to be counteracted by changes downstream in the transcription cycle, particularly at the promoter-proximal pause. We thus observed an increase in the release of polymerases from the promoter-proximal pause and a change in elongation characteristics when the cellular levels of RNAPII are lowered. It is important to stress that initiation frequency fundamentally restricts the number of full-length RNAs produced; a decrease in initiation rates can only be compensated for by an increase in RNAPII processivity across genes, and not, for example, by faster transcript elongation^63^. Indeed, a decrease in initiation rates can be compensated for by changes in promoter-proximal pausing *only* if the promoter-proximally paused polymerase can block or restrict the association of a new, initiating polymerases^63–65^, or if premature termination of transcription at the promoter-proximal transcription checkpoint is reduced, such as via decreased Integrator-mediated dissociation of early elongation complexes. While the precise mechanisms underlying these intriguing connections merits future investigation, we note that Integrator-and NELF-RNAPII interactions are indeed decreased as RNAPII dosage is lowered. Moreover, the pool of active P-TEFb and its interaction with RNAPII increase.

Together, these changes not only indicate intricate mechanistic connections between RNAPII dosage and the basic properties of transcription but also shed new light on the complex, dynamic interplay between transcription initiation, promoter-proximal pausing, and pause release.

### mRNA splicing and transcription termination

Arguably, one of the most surprising results of our study is that lowering RNAPII dosage so dramatically affects both mRNA splicing and transcription termination. While the precise mechanisms underlying these effects remain to be established, mRNA splicing and 3’ end processing^44,66^ are co-transcriptional processes known to be tightly linked to RNAPII elongation, and several models of transcription termination posit that slowing down of the polymerase at the 3’ end of genes is required for efficient transcription termination^18^. Indeed, an RPB1 point-mutation causing fast RNAPII elongation also results in a termination delay, i.e. readthrough^67^. Intriguingly, transcription readthrough has also been observed in several different stress conditions, including after heat shock^68,69^, hyperosmotic stress^70^, oxidative stress^71^, hypoxia^72^, viral infections^73^, and even in cancer^74^. The mechanisms underlying transcription readthrough (termination defects) in these conditions are still unclear, but the observation that relatively modest reductions in RNAPII abundance bring about alterations in both the elongation rate of the remaining RNAPII molecules, as well as increased transcription readthrough, raises the intriguing possibility that fluctuations in RNAPII levels might somehow contribute to the production of ‘downstream-of-genes’ transcripts^75^ observed in different stress conditions.

We also discovered that alternative mRNA splicing is highly sensitive to RPB1 dosage. While the number of significant alternative splicing (AS) events increased with moderate RPB1 depletion, this effect plateaued beyond a ∼50% reduction in RPB1 levels.

Our data thus show that splicing in specific subsets of genes is remarkably sensitive to changes in RNAPII levels, with their splicing decisions becoming increasingly biased as RNAPII levels decline. Interestingly, several of the AS events induced by depletion of RPB1 were also observed in cells expressing either fast or slow elongating RNAPII mutants^76^ (data not shown), supporting the idea that transcription kinetics play a central role in splice site selection when RNAPII levels are lowered. Altered expression of spliceosome components and snRNAs in response to RPB1 depletion may further shape the splicing landscape, either by changing the availability of specific regulators or by changing the efficiency or timing of spliceosome assembly. Finally, it is worth noting that many of the alternatively spliced genes play a role in cell cycle regulation, suggesting that splicing modulation might serve as an additional layer of control over cell cycle progression in response to changes in RNAPII abundance.

### RPB1 dosage sensitive genes

Given the global nature of the altered transcription dynamics observed upon reduced cellular levels of RNAPII, why then are some genes more sensitive to RPB1 dosage than others? Interestingly, dosage-sensitive genes are in general relatively short, and our data suggests that their expression is predominately regulated at the level of transcription initiation, rather than through promoter-proximal pausing or elongation. An extreme example of this is genes encoding snRNAs, which are only ∼200bp long. A mere 20% reduction in RPB1 levels greatly affected the recruitment of RNAPII to the promoters of snRNA genes, which might be explained at least partly by the fact that they possess a promoter structure distinct from that of protein-coding genes^77^. In contrast, RPB1 dosage-<u>in</u>sensitive genes are long, and their expression appears to be more heavily regulated at the promoter-proximal pause site than that of RPB1 dosage-sensitive genes. As such, the transcription of these dosage-insensitive genes benefits substantially from increased RNAPII release from the promoter proximal pause site in low RPB1 conditions, in effect buffering their expression to alterations in RNAPII dosage. Intriguingly, many general transcription initiation factor genes are part of this dosage-insensitive category. This raises the possibility that evolution has fine-tuned the regulation of these genes to render them less sensitive to changes in RNAPII levels. One obvious possibility is that their maintained expression allows them to help minimize the effect of suboptimal RNAPII levels on initiation. After all, if not only RNAPII, but also general transcription factors such as TFIID become limiting, one would expect exponential negative effects on transcription initiation.

### Implications for disease

The intriguing connection between RPB1 dosage (in)sensitive genes and biological function also has implications for disease. Cancer cells are often marked by hyper-transcription, a phenomenon associated with poor prognosis^78,79^. This genome-wide increase in transcriptional output is thought to result from increased activity of general transcription factors and co-activators^79–83^. Notably, however, overexpression of RNAPII subunits has also been linked to reduced survival rates in cancer patients^84^. Especially *POLR2A* is frequently found upregulated in tumor cells compared to normal tissues^85–87^. Our findings suggest that a slight increase in RPB1 levels leads to a mild, genome-wide elevation of transcription activity, raising the possibility that increased abundance of RPB1 may contribute to cancer-associated hyper-transcription or at least alterations to the gene expression characteristics of the cell. In apparent agreement with this, mutations in the *ARMC5* gene encoding the substrate recognition subunit of the RPB1-directed E3 ubiquitin ligase, CRL3^ARMC^^5^ results in primary bilateral macronodular adrenal hyperplasia, correlating with a significant increase in RPB1 levels and altered gene expression in adrenal cells^49^. Moreover, it is noteworthy that occupancy of RNAPII on replication-dependent histone genes, which are the genes most sensitive to RPB1 dosage (Figure 2G; Supplemental Figure 7C), has recently been found to correlate with tumor aggressiveness^88^. This opens the intriguing possibility that targeted modulation of RPB1 levels holds therapeutic potential, which might warrant further investigation. Taken together, these results highlight a compelling link between RPB1 dosage, gene regulation, and disease.

Multiple individuals have been identified with *de novo* heterozygous variants of likely pathogenic origin in the *POLR2A/RPB1* gene^19,20,89,90^. Several of these variants result in in-frame deletions, or truncation of the RPB1 protein^19,20^. As such, even though the effect of RPB1 protein levels of these pathogenic variants has not been investigated, the broad-spectrum developmental symptoms in individuals with these variants are thought to result from haploinsufficiency^19^. This idea is further supported by our data, which demonstrates that cells show a dramatic response even to less than 50% RPB1 loss. Even though the dTAG degradation system used here obviously does not reproduce the complex regulatory processes that occur during human development or maintain the feedback loop which allows upregulation of the remaining *RPB1* allele when RPB1 protein levels decrease, our results provide a putative disease mechanism underlying the symptoms observed in individuals with pathogenic variants in *POLR2A*.

## Acknowledgements

This work was supported by a Laureate grant from the Novo Nordisk Foundation (NNF19OC0055875) and a Center of Excellence grant (DNRF166) from the Danish National Research Foundation to J.Q.S., and by EMBO Postdoctoral Fellowships ALTF 2021-498 (to A.M.O.), ALTF 2020-260 (to M.N.G.), ALTF 2020-911 (to N.N.M.) and European Union’s Horizon Europe Marie Sklodowska-Curie grant 101066734 (to A.M.O.) and 101023584 (to M.N.G.). It was also funded by the Francis Crick Institute (FCI), which receives funding from Cancer Research UK [FC001166], the UK Medical Research Council [FC001166], and the Wellcome Trust [FC001166]. We would like to thank Jens Vilstrup Johansen for help with bioinformatics analysis, and members of the Svejstrup lab and CGEN for insightful discussions. The Crick Advanced Sequencing Facility and the CPR/ReNEW/ICMM Genomics Platform are thanked for support with sequencing and help with High-Performance Computing. We also acknowledge the Core Facility for Integrated BioImaging and the Flow Cytometry & Single Cell Core Facility at the University of Copenhagen (UCPH). Mass spectrometry-based proteomic analyses were performed by the Proteomics Research Infrastructure (PRI) at the UCPH, supported by the Novo Nordisk Foundation (NNF) (NNF19SA0059305).

## Data availability

The mass spectrometry proteomics data have been deposited to the ProteomeXchange Consortium via the PRIDE partner repository with the data set identifier PXD068019 (https://www.ebi.ac.uk/pride/archive/projects/PXD068019).

Genome-wide data has been deposited in the NCBI Gene Expression Omnibus repository with accession codes GSE307865, GSE307864, GSE307860, GSE307861 and GSE307769.

## Methods

### Cell lines and culture conditions

Flp-In T-REx HEK293 cells (Thermo Fisher Scientific), as well as RPB1 degron cell lines, were cultured at 37 °C/5% CO_2_ in standard Dulbecco’s modified Eagle’s medium (DMEM) supplemented with 10% FBS, 100 U/mL penicillin and 100 μg/mL streptomycin. RPB1 degron cells expressing the RPB1 transgene under a doxycycline-inducible promoter were cultured in standard Dulbecco’s modified Eagle’s medium (DMEM) supplemented with 10% tetracycline-free FBS, 100 U/mL penicillin and 100 μg/mL streptomycin. To induce expression of the transgene, varying concentrations of doxycycline were added to the medium for 48-72 hours prior to harvest.

### Generation of plasmids

The plasmid containing sgRNA targeting the N-terminus of *POLR2A* (NOE64-pX330A-sgPOLR2A-sgPITCh) was generated as follows. Top ranking sgRNAs targeting human *POLR2A* were identified using CHOPTOP, then hits targeting the N-term of the gene and closest to the first codon were selected. Sense and antisense oligos for the sgRNA (Table S1) were annealed by mixing 100pmol of each oligo in 1X annealing buffer (50mM KCl, 5mM MgCl_2_, 25 mM Tris-Cl pH 7.5), followed by incubation at 95 °C for 5 minutes and slow cool down to room temperature overnight. Annealed primers were then diluted 1:100 in 100mM Tris pH 7.5 and assembled into a BbsI-cut Cas9 vector GW223-pX330A-sgX-sgPITCh at a 1:3 vector:insert molar ratio using NEBuilder 2× HiFi assembly (New England Biolabs).

For generation of the 3x linker donor plasmid (NOE63-pCRIS-PITChv2-Puro-dTAG-3x-POLR2A-N), the FKBP12^F36V^ degron tag, puromycin resistance cassette and HA tags, were PCR amplified from the GW212-pCRIS-PITChv2-Puro-dTAG-BRD4N with primers (Table S1) containing 22bp microhomology arms corresponding to the genomic sequences flanking the POLR2A sgRNA cleavage site. The resulting linear fragment was then assembled into the MluI-linearized GW212-pCRIS-PITChv2-Puro-dTAG-BRD4N plasmid using NEBuilder 2× HiFi assembly (New England Biolabs).

To obtain donor templates with variable linker sizes, we obtained a synthetic DNA sequence containing 15x GGGS repeats from Genscript. Fragments of variable linker length were PCR amplified from the synthetic DNA template using a non-specific reverse primer that recognized the repetitive 3x linker. The randomly amplified 3x, 6x, 9x, 12x and 15x repeats were then assembled using NEBuilder 2× HiFi assembly into a PCR-linearized vector derived from the original 3x linker donor plasmid described above (NOE63-pCRIS-PITChv2-Puro-dTAG-3x-POLR2A-N). The resulting plasmids were screened using Sanger sequencing until the correct sequence was obtained. The 3x-mEGFP linker fragment was PCR amplified from a pKK-mEGFP-PS-TEV-3xFLAG plasmid and similarly cloned into the PCR-linearized vector derived from the original 3x linker donor plasmid. The 6x linker and 3x-mEGFP linker degron plasmids were then further subcloned to remove the hygromycin cassette, to facilitate recombination of rescue plasmids into the FRT locus.

To generate a plasmid expressing the RPB1-His coding sequence under the promoter of the human *UBC* gene, a pFRT-TO-wtRPB1-His-siR plasmid was digested with MluI and NotI to remove the inducible CMV promoter. A fragment containing 4kb genomic sequence upstream of the start codon of the human *UBC* gene was then PCR amplified from genomic DNA and cloned into the linear backbone using NEBuilder 2× HiFi assembly (New England Biolabs).

### Generation of stable cell lines

To generate POLR2A degron cell lines, Flp-In TREX HEK293 cells were seeded in 6-well plates in antibiotic-free media and transfected with 2μg cutting plasmid (NOE64-pX330A-sgPOLR2A-sgPITCh) and 1μg donor plasmid using lipofectamine 2000. After four days, cells were transferred to 10cm plates and 2.5ug/mL puromycin was added to the medium to select for edited cells. Cells were then diluted to single cells using limiting dilution techniques and grown to confluency. Genomic DNA from single clones was extracted using 30uL of DirectPCR Lysis Reagent supplemented with 400ug/mL proteinase K and incubation overnight at 55 °C. Proteinase K was then inactivated by incubating samples at 95 °C for 20 minutes, and 3uL of DNA was used for genotyping with primers that specifically hybridize outside the insert region (Table S1). PCR reactions were run on a 1% agarose gel to test for homozygosity and sent for Sanger sequencing to confirm the genotype.

Stable cell lines overexpressing RPB1 were generated by integration of the pFRT-TO-wtRPB1-FLAG-siR and pFRT-*UBC*-wtRPB1-His-siR constructs into the FRT locus of the RPB1 degron cell lines. Briefly, cells were seeded in antibiotic-free media in 6-well plates and transfected with a 9:1 mass ratio of RPB1 plasmid and pOG44 plasmid using Lipofectamine 3000. After 48hrs, cells were split to a 10cm plate containing selection medium (1X DMEM + 10% FBS + 100ug/mL hygromycin + 15ug/mL blasticidin + 1% Penn/Strep). The selection medium was replaced every 2-3 days until the formation of colonies was apparent. Individual colonies were then picked and expanded in 24-well plates while maintaining them in selection medium. Finally, expression of the RPB1 transgene was confirmed using flow cytometry and western blot.

### Cell growth analysis

For cell growth assays, 10k cells per dTAG^V^-1 concentration were seeded in triplicate in a 96-well plate. The following day, the medium was replaced with medium containing different concentrations of dTAG^V^-1. The growth of these cells was then monitored using an Incucyte S3 Live-Cell Analysis System (Sartorius). Images of live cells were captured every 4 hours for up to five days and confluency was calculated using the Incucyte Classic Confluence Analysis workflow.

### Hybridization Chain Reaction (HCR)-based fluorescent in situ hybridization (FISH)

Cells were seeded at a density of 30,000 cells per well in 8-well microscopy chamber slides pre-coated with poly-D-lysine. On the following day, cells were treated with different concentrations dTAG^V^-1 or DMSO. After sixteen hours, cells were washed once with 1× PBS and fixed with 4% formaldehyde for 10 min at room temperature. Following fixation, cells were washed three times with 20 mM Tris-HCl (pH 7.5) in PBS for 5 min each, and then permeabilized with 70% ethanol for 1 h at room temperature. Afterwards, cells were washed twice with 2× SSC and pre-hybridized with 30% probe hybridization buffer (30% formamide, 5X sodium chloride sodium citrate (SSC), 9mM citric acid (pH6.0), 0.1% Tween-20, 50ug/mL heparin, 1X Denhardt’s solution, 10% dextran sulfate) for 30 min at 37 °C. HCR probe solutions were prepared by combining HCR split initiator probes (20nM final concentration; 5 probe pairs for *POLR2A*; 1 probe pair for oligodT; 1 probe pair for scrambled (Table S1)) in pre-warmed 30% probe hybridization buffer and incubating at 37 °C for 30 min. Pre-hybridization buffer was removed and replaced with the pre-made HCR probe solution, and samples were incubated overnight at 37 °C. The following day, excess probes were removed by washing four times for 5 min with 30% probe wash buffer (30% formamide, 5X SSC, 9mM citric acid (pH6.0), 0.1% Tween, 50ug/mL heparin) at 37 °C, followed by two washes with 5× SSCT (5X SSC + 0.1% Tween-20) at room temperature. Cells were then incubated in amplification buffer (5X SSC, 0.1% Tween-20, 10% dextran sulfate) for 30 min at room temperature. Fluorescently labeled hairpins were snap-cooled by heating at 95 °C for 90 s and cooling to room temperature for 30 min before being added to the amplification buffer (30nM final concentration). Cells were then incubated with the hairpin solution overnight at room temperature. On the final day, samples were washed three times with 5× SSCT at room temperature to remove excess hairpins. Cells were blocked with 1 mL of 4× SSC containing 3% BSA for 30 min at room temperature, followed by incubation with rabbit anti-DIG primary antibody (clone D8Q9J, Cell Signaling Technology, 1:500 dilution in 4× SSC + 3% BSA) for 1 h at room temperature. After three washes with 4× SSC + 0.1% Triton X-100, cells were incubated with donkey anti-rabbit secondary antibody conjugated to Alexa Fluor 488 (1:500 dilution in 4× SSC + 3% BSA) for 1 h at room temperature in the dark. Samples were then washed again three times with 4× SSC + 0.1% Triton X-100. Nuclei were counterstained with DAPI (1:5000 dilution in 1X PBS) for 10 min at room temperature, followed by a final PBS wash and mounting of a coverslip with ProLong Diamond Antifade Mountant. Slides were imaged using a Zeiss LSM 980 laser scanning confocal microscope.

### Immunofluorescence

20,000 cells were seeded in the 8-well Nunc Lab-Tek II CC^2^ Chamber Slide system and dTAG^V^-1 was added to the medium the following day. After 24 hours, cells were fixed for 15 minutes with 4% formaldehyde, followed by three 1X PBS washes and permeabilization for 10 minutes with 0.25% Triton-X/PBS. After blocking for 1 hour with 3% BSA/PBS, cells were incubated with primary antibodies (diluted 1:300 in 3% BSA/PBS) overnight at 4 °C. Cells were then washed 4 times with 1X PBS, and incubated with secondary antibody (diluted 1:500 in 3% BSA/PBS) for 1 hour at RT. Afterwards, cells were washed 4 times with 1X PBS and nuclei were stained with DAPI (diluted 1:5000 in 1X PBS) for 15min at RT.

Coverslips were then mounted on the slide with Prolong Diamond Antifade and cells were imaged using a Zeiss LSM 710 laser scanning confocal microscope.

### Flow cytometry

Cells were seeded in 6-well plates and treated with different concentrations of dTAG^V^-1 the following day. For cell cycle analysis, cells were pulsed with 10uM EdU for 30 minutes prior to harvest. To evaluate transcription, cells were pulsed with 500uM EU for 1 hour prior to harvest. 16hrs after addition of dTAG^V^-1, cells were trypsinized and washed once with 1% BSA/PBS. They were then fixed for 10 minutes at RT with 4% formaldehyde, washed with 1% BSA/PBS, and barcoded for 30 minutes at RT with Dylight 488 and 800 NHS esters dissolved in 70% ethanol (serial dilutions ranging from 0-5ug/mL). After two washes with 1% BSA/PBS, samples were pooled into a single tube and ∼1-2 million cells were incubated with primary antibodies (diluted 1:300 in 1% BSA/PBS) overnight at 4 °C. Cells were then washed once with 1% BSA/PBS, and unstained, but permeabilized cells were added to the tube prior to incubation with secondary antibodies (diluted 1:500 in 1% BSA/PBS) for 1 hour at RT. Afterwards, cells were washed once with 1% BSA/PBS and stained with DAPI (diluted 1:5000 in 1% BSA/PBS) for 15 minutes at RT. Cells pulsed with EU or EdU were further incubated with the Click-iT Plus reaction cocktail for 30 minutes at RT (Click-iT Plus EdU Alexa Fluor 647 Flow Cytometry Kit, #C10634), followed by a wash with 1% BSA/PBS. Samples were then run on the FACSymphony A5 and deconstructed into the individual dTAG^V^-1 conditions using the AF488 and IR800 signal. Protein levels were further analyzed using FlowJo software, including export of median fluorescent values and subtraction of background fluorescence from unstained controls. Protein expression was then normalized to DMSO conditions to facilitate averaging across a minimum of three independent experiments. Statistical significance was determined using GraphPad Prism, through a mixed-effects analysis with Geisser-Greenhouse correction, followed by Dunnett’s multiple comparisons test.

### Western blotting

For whole cell extracts, cell pellets were collected in ice-cold 1X PBS and resuspended in RIPA buffer (150mM NaCl, 50mM Tris-HCl pH8.0, 1% Triton X-100, 0.5% sodium deoxycholate, 0.1% SDS), supplemented with protease inhibitors, phosphatase inhibitors, and 1:1000 BaseMuncher endonuclease. Samples were incubated on ice for 15 minutes, followed by centrifugation at 21,000xg for 15 minutes. Protein concentration of the clarified extracts was determined using the Pierce™ Rapid Gold BCA Protein Assay Kit. Laemmli with 50mM DTT was then added to the cell lysates and samples were boiled at 98 °C for 5 minutes. 30ug protein per sample was loaded on 4%–15% TGX gels (Bio-Rad, 56711084/5) and separated by SDS-PAGE for 45 minutes at 180V. Samples were then transferred to nitrocellulose membranes (GE Healthcare Life Sciences, 10600002) using the Biorad Trans-Blot Semi-Dry Transfer cell. Equal loading of protein mass was confirmed by Ponceau S staining, after which membranes were blocked in Intercept (TBS) Blocking Buffer (Licor) for one hour at RT, and incubated with primary antibodies (1:1000 in Licor Intercept (TBS) Blocking Buffer + 0.1% Tween-20 + 0.02% sodium azide) overnight at 4 °C. Membranes were then washed 3 times with TBST and incubated with secondary antibodies (1:2000 in Licor Intercept (TBS) Blocking Buffer + 0.1% Tween-20) for 1 hour at RT. Afterwards, membranes were washed 3 times with TBST, and fluorescent signal was detected using an Amersham Typhoon fluorescence imaging system. Quantitative analysis was performed using Image J.

### qRT-PCR

Cells were seeded in 24-well plates, followed by addition of dTAG^V^-1 the day after. After 16 hours, cells were harvested in 200uL TRIzol, and RNA was extracted using the RNeasy Mini Kit (Qiagen, #74106) as per the manufacturer’s instructions. RNA concentration was measured using Nanodrop and 1ug total RNA was reverse transcribed into cDNA using random hexamers. To control for the presence of genomic DNA contamination, a parallel reaction was set up for each sample without reverse transcriptase. cDNA was diluted 50-fold and used as template for qRT-PCR using gene-specific primers (Table S1) and iTaq Universal SYBR Green supermix (Biorad, #172-5124).

### Immunoprecipitation

Cells were seeded in 15cm plates, followed by treatment with either DMSO, 32nM dTAG^V^-1, 62nM dTAG^V^-1, or 125 nM dTAG^V^-1 for 16 hours, and harvest in ice-cold 1x PBS. Cells were fractionated by initially lysing cells on ice for 20 minutes in a lysis buffer (20mM HEPES pH 7.5, 150 mM NaCl, 2.5 mM MgCl_2_, 0.15% (v/v) NP-40 alternative, 10% (v/v) glycerol, 1mM DTT, 1x protease inhibitor, 1x phosphatase inhibitor). To isolate nuclei, the cell lysate was placed on top of a sucrose cushion (20mM HEPES pH 7.5, 150 mM NaCl, sucrose 25 % (v/v), 10% (v/v) glycerol, 1mM DTT, 1x protease inhibitor, 1x phosphatase inhibitor) and centrifuged at 1000g and 4°C for 5 minutes. Supernatant containing the cytosolic fraction was removed and the nuclear pellet was resuspended in chromatin digestion buffer (20mM HEPES pH 7.5, 150 mM NaCl, 1 mM MgCl_2_, 0.05% (v/v) NP-40 alternative, 10% (v/v) glycerol, 500 U/mL BaseMuncher, 1x protease inhibitor, 1x phosphatase inhibitor) and was incubated for 1h at 4°C on a rotating wheel. The nuclear chromatin extract was centrifuged at max speed at 4°C for 10 minutes. The supernatant was removed and saved as nuclear extract. To extract the tightly chromatin-bound proteins, the pellet was further resuspended in high salt chromatin digestion buffer (20mM HEPES pH 7.5, 300 mM NaCl, 1 mM MgCl_2_, 0.05% (v/v) NP-40 alternative, 10% (v/v) glycerol, 500 U/mL BaseMuncher, 1x protease inhibitor, 1x phosphatase inhibitor) and incubated for 1h at 4°C on a rotating wheel. The high salt chromatin extract was hereafter diluted with 1 volume dilution buffer (20mM HEPES pH 7.5, 1 mM MgCl_2_, 0.05% (v/v) NP-40 alternative, 10% (v/v) glycerol, 1x protease inhibitor, 1x phosphatase inhibitor). The low salt and high salt nuclear extracts were pooled and centrifuged at max speed at 4°C for 5 minutes. Protein concentration of the clarified extracts was determined using the Pierce 660 protein kit assay and diluted to equal concentrations, so the total mass of protein for input was 800 ug. Lysates for each condition were split into technical replicates (3 per antibody) and incubated overnight at 4°C with protein G-coupled mouse anti-RNAPII CTD (4H8) or mouse anti-IgG antibodies (1µg antibody per 10µL protein G dynabeads). The following day, beads were washed three times in IP buffer (20mM HEPES pH 7.5, 150 mM NaCl, 1 mM MgCl_2_, 0.05% (v/v) NP-40 alternative, 10% (v/v) glycerol, 1x protease inhibitor, 1x phosphatase inhibitor). For mass spectrometry analysis, the beads were then further washed twice in ice-cold 1xTBS with 1x protease inhibitor and 1x phosphatase inhibitor. Following the last wash the beads were transferred to new 1.5 mL Eppendorf tubes.

### Mass spectrometry

Washed beads were incubated for 30 min with elution buffer 1 (2 M Urea, 50 mM Tris-HCl pH 7.5, 2 mM DTT, 20 μg/mL trypsin) followed by a second elution for 5 min with elution buffer 2 (2 M Urea, 50 mM Tris-HCl pH 7.5, 10 mM Chloroacetamide). Both eluates were combined and incubated at RT overnight. Tryptic peptide mixtures were acidified to 1% TFA and loaded on Evotips (Evosep). Peptides were separated on a 15 cm, 150 μM ID column packed with C18 beads (1.9 μm) (Pepsep) on an Evosep ONE HPLC applying the 30SPD method and injected via a CaptiveSpray source and 10 μm emitter into a timsTOF pro mass spectrometer (Bruker) operated in PASEF mode.

### Subcellular fractionation for pTEFb activity

Separation of large and free forms of P-TEFb by differential salt extraction was performed essentially as described previously^39^. Cell pellets were collected in ice-cold 1X PBS and were resuspended in 80 μL of Buffer A (10 mM KCl, 10 mM MgCl, 10 mM HEPES pH 8.0, 1 mM EDTA, 1 mM DTT, 1× protease inhibitor, 0.5% NP-40) and incubated on ice for 10 min. Nuclei were pelleted by centrifugation at 5,000 × g for 5 min at 4 °C. The supernatant, corresponding to the cytosolic extract (CE), was collected and mixed with an equal volume of 2× Laemmli buffer. This fraction contains the long, 7SK-bound (inactive) isoform of P-TEFb. The nuclear pellet was washed once with 200 μL of Buffer A and centrifuged again at 5,000[× g for 5 min. Nuclei were then resuspended in 80 μL of Buffer B (450 mM NaCl, 1.5 mM MgCl, 20 mM HEPES pH 8.0, 0.5 mM EDTA, 1 mM DTT, 1× protease inhibitor, 0.5% NP-40) and incubated on ice for 10 min. Nuclear debris was pelleted by centrifugation at maximum speed (≥20,000 × g) for 10 min at 4 °C. The supernatant, corresponding to the nuclear extract (NE), was collected and mixed with an equal volume of 2× Laemmli buffer. This fraction contains the free, active isoform of P-TEFb lacking 7SK association.

### TT_chem_-seq

TT_chem_-seq was performed as described previously^32^. Briefly, 3 million RPB1 degron cells and *UBC*-RPB1-His-siR overexpressing cells were seeded in triplicate in 10cm plates, and different concentration dTAG^V^-1 were added the next day. After 16 hours, 4SU was added directly to the cell culture medium to a final concentration of 1mM. After 15 minutes incubation, medium was aspirated, and the labelling was terminated by addition of 1mL Trizol to each plate. Total RNA was extracted using TRIzol/chloroform extraction, followed by isopropanol precipitation. RNA concentration was measured using Qubit RNA BR Assay Kit (Thermo Fisher, Q10210). For each sample, 1 μg of *S. cerevisiae* 4TU-labeled RNA was mixed with 100 μg of the purified 4SU labeled RNA in a total volume of 100μl made up with RNase-free water. RNA was then fragmented through the addition of 20uL 1M NaOH and incubation on ice for 20 minutes. Fragmentation was terminated through the addition of 80uL 1M Tris-HCl, pH6.8, followed by two rounds of clean-up with Micro Bio-Spin P-30 gel columns (Bio-Rad, 7326250). The fragmented RNA was biotinylated in a total volume of 250uL, containing 10 mM Tris-HCl pH 7.4, 1 mM EDTA and 5 mg MTSEA biotin-XX linker (Biotium, BT90066) for 30 min at room temperature in the dark. Afterwards, RNA was purified by phenol/chloroform extraction and isopropanol precipitation. The RNA was denatured at 65C for 10 minutes and incubated with 100uL μMACS Streptavidin beads for 15 minutes at RT, followed by purification using a μMACS Streptavidin Kit (Miltenyi, 130-074-101) according to the manufacturer’s instructions. The biotinylated RNA was further cleaned up using the RNeasy MinElute kit (QIAGEN, 74204) and quantified using Qubit RNA HS Assay Kit. Libraries were prepared at the Francis Crick Institute using the KAPA RNA HyperPrep Kit (Roche, 08098093702) with the fragmentation step omitted. The libraries were then sequenced on an Illumina NovaSeq 6000, yielding 50bp paired-end reads per sample.

### DRB/TT_chem_-seq

RPB1 degron cells were seeded at a density of 4 million cells in a 10 cm plate and were either treated with DMSO or different concentrations of dTAG^v^-1 for 16 hours before harvest. After 12.5 hours, transcription was synchronized by the addition of 100uM DRB (5,6-dichloro-1-β-D-ribofuranosylbenzimidazole) for 3.5 hours. DRB was then removed by three 1X PBS washes, facilitating the release of RNAPII from the promoter-proximal pause for 15, 30, or 45 minutes prior to harvest. During the final 10 minutes prior to harvest, cells were pulsed for 1mM 4-thiouridine, to metabolically label nascently transcribed RNA. Afterwards, medium was aspirated, and the labelling was terminated by addition of 1mL Trizol to each plate. Extraction of nascent RNA occurred as described for TT_chem_-seq and included yeast 4TU-labelled RNA for spike-in. Libraries were prepared from biotinylated RNA using the NEBNext Ultra II Directional RNA Library Prep Kit for Illumina (without additional fragmentation) and NEBNextMultiplex Oligos for Illumina (Unique Dual Index UMI adapters RNA Set 1) and were sequenced on an Illumina NextSeq 2000 (Genomics Platform at University of Copenhagen), yielding 50bp single-end reads.

### PRO-seq

PRO-seq was performed essentially as described previously^91^. Briefly, DMSO or dTAG^V^-1-treated cells or Drosophila S2 cells were permeabilized and counted. 1 million permeabilized RPB1 degron cells (nuclei) per sample were mixed with 5% spike-in Drosophila S2 permeabilized cells (nuclei) in 50 μl, then mixed with 50 μl 2X Run-On Master Mix containing biotin-tagged CTP and UTPs (PerkinElmer, NEL542001EA-NEL545001EA) to perform the run-on reaction at 37°C for 5 min. The reaction was terminated by adding TRIzol LS followed by total RNA extraction. Total RNA was denatured by adding NaOH to 200 mM and then purified on a Bio-Rad RNase free P-30 column following the manufacturer’s instructions. Then the 3’ RNA adaptor ligation was performed, and the biotin labeled RNA was enriched on Streptavidin beads. Next 5’ hydroxyl repair and de-capping reactions were performed on-bead followed by 5’ RNA adaptor ligation. After these enzymatic reactions, the Streptavidin beads with bound biotin labeled RNA were washed. Biotin labeled RNA was released from the Streptavidin beads using TRIzol/chloroform extraction followed by precipitation. Reverse transcription was performed off-bead to get cDNA. 1.54 μl out of the 20 μl reverse transcription reaction was used to do a test amplification (qRT-PCR) to determine the amplification cycle number, then full-scale amplification was performed with 15 cycles. PCR products were cleaned up and selected on SPRI beads for sequencing. The libraries were then sequenced on an Illumina NextSeq 2000 (Genomics Platform at University of Copenhagen), yielding 100bp single-end reads.

### mRNA-seq

RPB1 degron cells were seeded at a density of 4 million cells in a 10cm plate and were either treated with DMSO or different concentrations of dTAG^V^-1 for 16 hours. Cells were harvested by the addition of 1mL Trizol to each plate. To isolate total RNA, one-fifth of chloroform was added to lysates and shaken well, followed by centrifugation (12000g, 5 minutes at 4C). The upper aqueous phase was then transferred to MaXtract high-density phase-lock tubes, that were previously prepared according to manufacturer’s instructions, and an equal volume of chloroform/isoamyl alcohol (24:1) was added. Lysates were then centrifuged again (12000g, 5 minutes at 4C) and the upper aqueous phase was transferred to a new tube. The RNA was then precipitated through the addition of 1.1X volumes of 100% isopropanol, incubation at room temperature for 20 minutes and centrifugation (12000g, 20 minutes at 4C). RNA pellets were washed once with 85% (v/v) ethanol, and air dried at room temperature before resuspension in Ultrapure DNase/RNase-free distilled water. 2ug total RNA was used for oligo(dT) purification. Sequencing libraries were then constructed at the Francis Crick Institute using a KAPA mRNA HyperPrep Kit and sequenced on an Illumina NovaSeq 6000, yielding 100bp paired-end reads per sample.

### CUT&RUN

CUT&RUN was performed essentially as per manufacturer’s instructions (#86652, Cell Signaling). 5 million cells were seeded in 10cm plates per condition and treated with different concentrations of dTAG^v^-1. After 16 hours, cells were trypsinized and fixed for 2 minutes at room temperature with 0.1% formaldehyde. Fixing was stopped by addition of 250mM glycine for 5 minutes, and cells were washed once with 1X wash buffer supplemented with spermidine and protease inhibitor cocktail. Cells for each condition were then aliquoted into 100 thousand cells per Lo-bind microcentrifuge tube and incubated with activated concanavalin A beads for 5 minutes at room temperate. After addition of antibody binding buffer, cells were incubated overnight at 4C with rabbit anti-HA (4uL per replicate, 3 replicates per dTAG^v^-1 condition), rabbit anti-Ser5+2P (4uL per replicate, 3 replicates per dTAG^v^-1 condition), and rabbit anti-IgG (5uL per replicate, 3 replicates per dTAG^v^-1 condition) antibody. Binding of pAG-MNase and DNA digestion was performed as per manufacturer’s instructions and included the addition of 50pg spike-in DNA. To reverse crosslinks, the eluted DNA was incubated with agitation overnight at 65C in the presence of 0.1% SDS and 133ng/ug proteinase K. DNA was purified using DNA Purification Buffers and Spin Columns (#14209 Cell Signaling) and 40uL of each sample was used for library preparation with the NEBNext Ultra DNA Library Prep Kit for Illumina (#E7370L, NEB). Unique dual indices (#E7395S, NEB) were used to facilitate multiplexing and the detection of PCR duplicates. Finally, libraries were pooled and sequenced on an Illumina NextSeq 2000 (Genomics Platform at University of Copenhagen), yielding 50bp paired-end reads.

### Bioinformatics analysis Mass spectrometry analysis

Raw mass spectrometry data were analyzed with MaxQuant (v1.6.15.0). Peak lists were searched against the swine or bovine Uniprot FASTA databases combined with 262 common contaminants by the integrated Andromeda search engine. False discovery rate was set to 1 % for both peptides (minimum length of 7 amino acids) and proteins. “Match between runs” (MBR) was enabled with a Match time window of 0.7, and a Match ion mobility window of 0.05 min. Relative protein amounts were determined by the MaxLFQ algorithm with a minimum ratio count of two.

All statistical analysis was performed using in-house developed python code, based on the automated analysis pipeline of the Clinical Knowledge Graph^92^. Protein entries referring to potential contaminants, proteins identified by matches to the decoy reverse database, and proteins identified only by modified sites, were removed. LFQ intensity values were normalized by log2 transformation and proteins with less than 70% of valid values in at least one group were filtered out. The remaining missing values were imputed using the mixed imputation approach. With this method, we look at missing values in samples belonging to the same group and impute with k-nearest neighbors (kNN) if there is at least 60% of valid values in that group, for that protein. The remaining missing values are imputed with the MinProb method (random draws from a Gaussian distribution; width = 0.2 and downshift = 1.8)^93^. Differentially enriched proteins in each group comparison were identified by SAMR multiclass test with permutation-based FDR correction for multiple hypothesis (FDR < 0.01, s0 = 1, permutations = 250), followed by posthoc pairwise comparison unpaired t-tests using the same parameters and permutation-based FDR correction.

### TT_chem_-seq analysis

Paired-end reads were demultiplexed using bcl2fastq (v2.20.0.422) and resultant FASTQ files were quality checked using MultiQC (v1.11). Paired-end reads were aligned to the GRCh38 and sacCer3 genomes using hisat2 (v2.2.1) with options--no-mixed--no-discordant-p 16--rna-strandness RF, followed by identification of PCR duplicates using picard Markduplicates (v2.27.3). For external reference normalization using yeast spike-in material, reads mapping to genes in the sacCer3 genome were counted using FeatureCounts from the subread package (v2.0.6; featureCounts-O-M-T 4-s 2--raction-t gene). The counts matrix was then passed to DESeq2’s estimateSizeFactors function and the resulting yeast sizeFactors were used as scaling factors to account for global changes in transcription.

BAM files were converted to strand-specific bigWig files using the bamCoverage function from the deeptools package (v3.5.5, options:--samFlagInclude 64--normalizeUsing None--scaleFactor ${scalefactor}-p 16--binSize 1--filterRNAstrand=forward/reverse). For visualization of single genes, signal of individual replicates was averaged using the wiggletools write_bg function (option: mean), followed by the bedGraphToBigWig tool from kentUtils. For metagene analyses, bigWig files of individual replicates were used to generate count matrices using the deeptools function computeMatrix scale-regions (v3.5.5, options:--missingDataAsZero-m 15000-a 5000-b 5000--binSize 100--transcriptID gene--transcript_id_designator gene_id-p 4). Profiles of the average signal were then generated using custom R scripts (v4.2.1).

To perform differential gene expression analysis, we first counted reads aligning to genes in GRCh38 genome (Ensembl release 109) using FeatureCounts from the subread package (v2.0.6; featureCounts-O-M-T 4-s 2--fraction-t gene). Differential gene expression was then determined with DESeq2 (alpha=0.01), using the yeast sizeFactors. For visualization of the data in MA-plots, the log2 fold-change was shrunk using the apeglm method.

### Elongation index calculation

The number of TT_chem_-seq reads in individual replicates mapping to the beginning (TSS to +1kb) and end (-1kb to TES) of non-overlapping protein-coding genes between 60-300kb in length was calculated using bedtools intersect (v2.31.0; options:-wb-bed-split-S). To calculate the elongation index, the number of reads at the end of genes were divided by the number of reads at the beginning of genes and averaged across the biological replicates.

### DRB/TT_chem_-seq analysis

Single-end reads were demultiplexed using bcl2fastq (v2.20.0.422) and resultant FASTQ files were quality checked using MultiQC (v1.11). Reads were aligned to the GRCh38 genome using hisat2 (v2.2.1) with default options, followed by identification of PCR duplicates using picard Markduplicates (v2.27.3). Further analysis was conducted as in Gregersen et al. (2020)^32^ using custom R scripts (v4.3.1). Briefly, bp-resolution read-depth coverage was calculated over intervals representing non-overlapping, protein-coding genes at least 60kb in width, extended-2kb:+120kb around the TSS region. Counts were normalized for sequencing depth (RPM) and meta-profiles were created by taking a trimmed mean across genes at each bp location. A smoothing spline was fitted to each meta-profile using R’s smooth.spline function (spar=0.3). Wave peaks were subsequently calculated as the maximum point on the spline for each sample, ensuring that wave peaks advanced with time. Elongation rates in kb/min were estimated by fitting a linear model to the wave peak positions of all samples (i.e. 15-, 30-, and 45-minutes post release) as a function of time.

### mRNA-seq analysis

Paired-end reads were aligned to the GRCh38 genome using hisat2 (v2.2.1) with options--no-mixed--no-discordant-p 16--rna-strandness RF, followed by identification of PCR duplicates using picard Markduplicates (v2.27.3). To perform differential gene expression analysis, reads mapping to genes in the GRCh38 genome (Ensembl release 109) were counted using FeatureCounts from the subread package (v2.0.6; featureCounts-O-M-T 4-s 2--fraction-t exon). Differential gene expression was then determined with DESeq2 (alpha=0.01).

### mRNA half-life calculation

Expression of protein-coding genes in the TT_chem_-seq and mRNA-seq datasets was calculated by converting gene and exon counts respectively (obtained from FeatureCounts) to TPM values. Genes with a basemean below 100 in the mRNA-seq data were filtered out. To infer mRNA half-lives, the stable mRNA expression values were then divided by the expression values of the nascent RNA for individual replicates and finally averaged for each condition.

### Alternative splicing detection

Alternative splicing events were detected in the mRNA-seq data using the Bioconductor package EventPointer^94^. To this end, paired-end reads were aligned to the GRCh38 genome using STAR (v2.7.11b; options:--outSAMattributes XS), followed by the generation of BAM files using samtools (v1.15.1). A splice graph constructed from v109 of the Ensembl human transcriptome and the aligned reads from all BAM files was then used to detect and quantify a range of alternative splicing events. Events were quantified and a differential detection analysis was conducted between the DMSO and each of the dTAG^V^-1 concentration samples. AFE/ALE and A3SS/A5SS annotations were swapped for genes on the negative strand for consistency. To determine whether alternative splicing events would affect the coding sequence or UTR of a gene, coordinates of the AS events were compared with those of the CDS/UTR of transcripts with support level 1 using bedtools intersect (v2.31.0; option:-s).

### CUT&RUN analysis

Paired-end reads were aligned against the GRCh38 genome with hisat2 (v2.2.1) using the--no-spliced-alignment,--no-mixed and--no-discordant options. Afterwards, multi-mappers were removed using samtools (v1.15.1), followed by removal of PCR duplicates using the UMI-tools dedup function (v1.1.4). BAM files were then converted to bigwig files for visualization using the deeptools (v3.5.5) bamCoverage function with options--normalizeUsing CPM,--binSize 10, and--extendReads. Normalization of RNAPII coverage to IgG, as well as normalization of Ser5p+Ser2p RNAPII coverage to total RNAPII was performed using the deeptools function bigwigcompare. Finally, matrices for metagene plots were generated using the deeptools computeMatrix function, followed by visualization of the profiles using custom R scripts.

### Analysis of PRO-seq

Adapters were trimmed from single-end reads with fastp using options -- disable_trim_poly_g,--disable_quality_filtering,--length_required 16,--length_limit 125,-a TGGAATTCTCGGGTGCCAAGGAACTCCAGTCAC. UMI sequences were then extracted from the fastq files using umi_tools extract. The resulting reads were aligned against a combined human/drosophila genome (GRCh38/dm6) using hisat2 (default settings) and PCR duplicates were subsequently removed using the umi-tools dedup function with default settings. BAM files were then converted to strand-specific bigwig files containing only the first nucleotide of the 5’ end of aligned reads, which corresponds to the 3’ end of RNA transcripts, using deeptools bamcoverage with options--normalizeUsing CPM-p 16--binSize 1--ilterRNAstrand=forward/reverse--Offset 1.

### Pausing index calculation

The number of PRO-seq reads in individual replicates mapping to promoter regions (-50nt to +300nt) and gene body (+300nt to TES) were calculated for all non-overlapping protein-coding genes longer than 2kb, using bedtools coverage (v2.31.0; option:-S). To calculate the pausing index, the number of reads in the promoter and gene body were first normalized to the length of each region and then divided by each other (normalized promoter coverage / normalized gene body coverage).

### Gene ontology analysis

Ensembl IDs of genes in each RPB1 dosage sensitivity bin were used for gene ontology analysis using the g:Profiler webserver. Driver terms for the resulting GO molecular functions, biological processes and cellular components were reported in Supplementary Figure S5.

### Motif enrichment

To identify motifs differentially represented in the promoter of RPB1 dosage sensitive and insensitive genes, the promoter sequence of non-overlapping genes expressed above 1 TPM (-100nt to +100nt) was extracted using bedtools getfasta (v2.31.0; options:-s-name). Enriched motifs were identified using the findMotifs.pl function (option:-len 6,8,10) from HOMER (v5.1) using genes downregulated from 30% depletion onwards as the foreground and the unaffected genes as the background.

## Supplemental figures and legends

**Figure S1.**
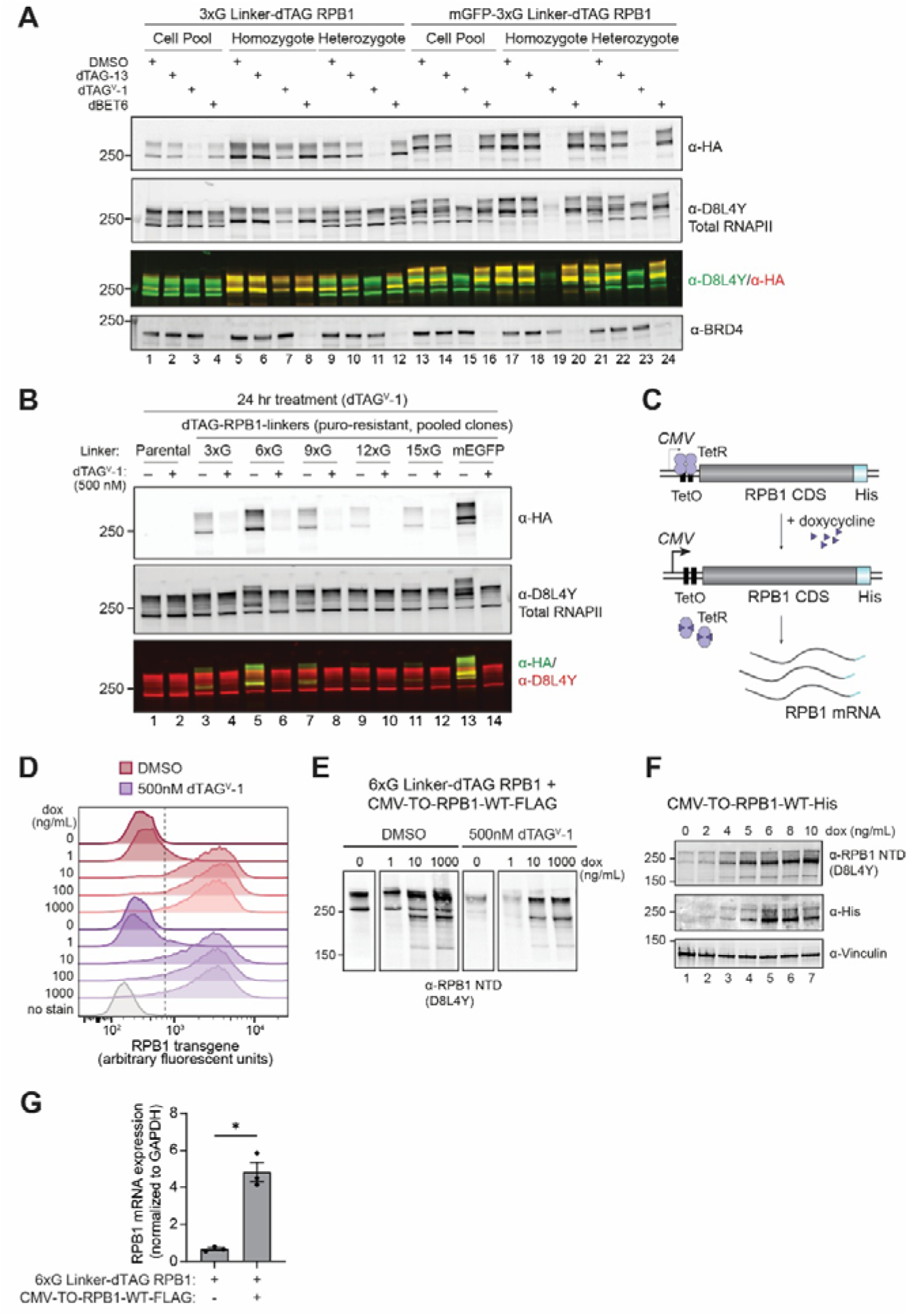
Related to. **Figure 1. (A)** Western blot analysis of RPB1 expression in the 3XG linker and mGFP-3XG linker RPB1 degron cell lines following 24 hour treatment with 500nM of different dTAG ligands. RPB1 degradation was evaluated using an antibody against the N-terminus of RPB1 and an anti-HA tag antibody. To evaluate the activity of the CRBN ubiquitin ligase, which is recruited by dTAG-13, treatment with dBET6 was included, which recruits the CRBN ubiquitin ligase to degrade BRD4 **(B)** Western blot analysis of RPB1 expression in cells expressing RPB1 degron construct with different linker size following 24 hour treatment with 500nM dTAG^V^-1. RPB1 degradation was evaluated using an antibody against the N-terminus of RPB1 and an anti-HA tag antibody. **(C)** Schematic of the switchover system, whereby an RPB1 transgene is expressed from a doxycycline-inducible promoter. **(D)** Flow cytometry analysis of the RPB1 transgene expression following 48hr induction with different doxycycline concentrations and treatment with dTAG^V^-1. RPB1 transgene expression was determined using an antibody against the His-tag. **(E-F)** Western blot analysis of total RPB1 expression following 48hr induction with different doxycycline concentrations and treatment with dTAG^V^-1. RPB1 expression was determined using an antibody against the N-terminus of RPB1. **(G)** qRT-PCR results for the relative expression of *POLR2A* mRNA in the 6XG linker RPB1 degron cell line and the 6XG linker RPB1 degron cell line with stable integration of the doxycycline-inducible RPB1 transgene following 48 hours treatment with 1ug/mL doxycycline. *POLR2A* expression is normalized to *GAPDH.* Statistical significance was determined by a Student’s t-test. *=P-value<0.05.

**Figure S2.**
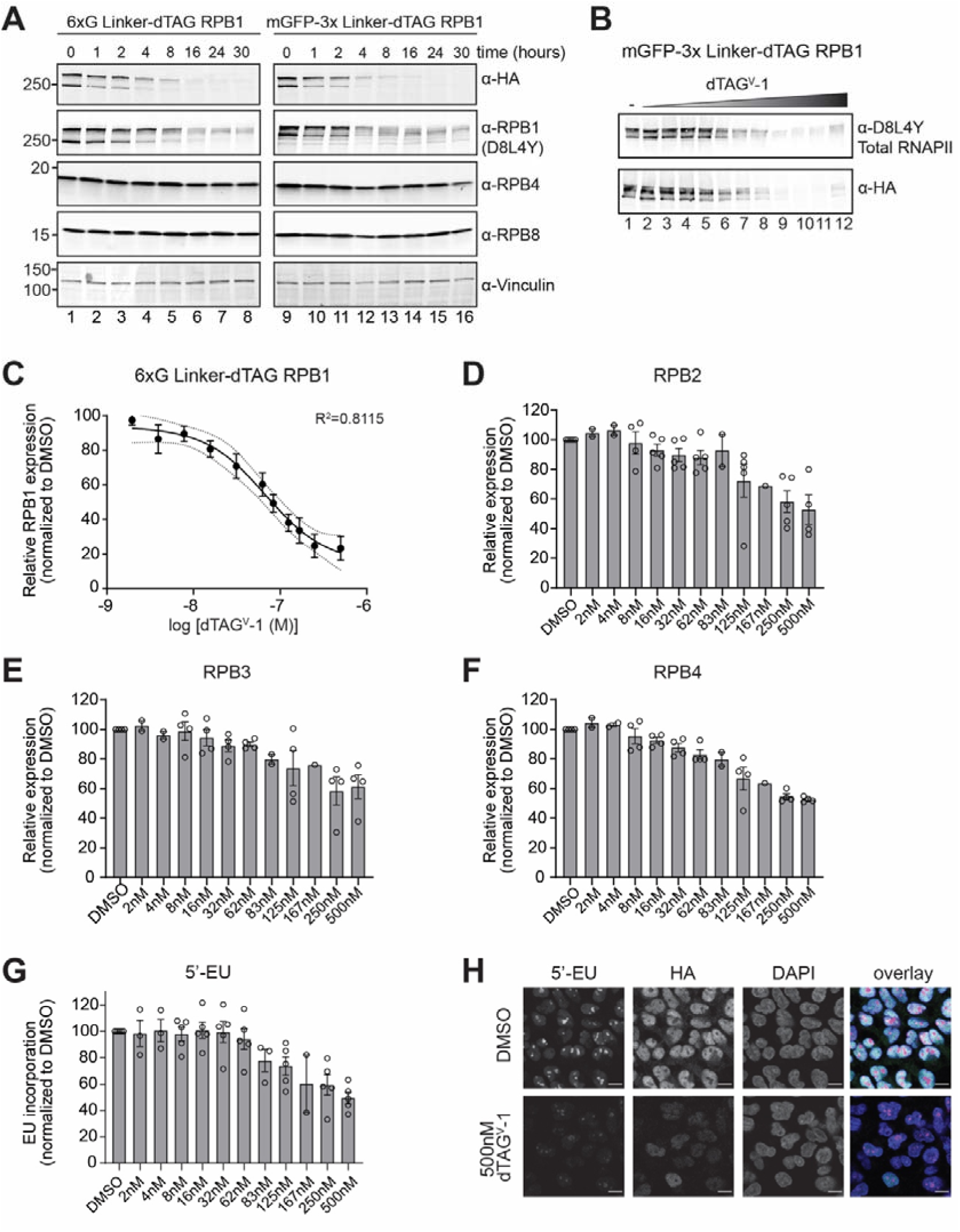
Related to. **Figure 1. (A)** Western blot analysis of RNAPII subunit expression in the 6XG linker and mGFP-3XG linker RPB1 degron cell lines harvested at different times following 500nM dTAG^V^-1 treatment. **(B)** Western blot analysis of dose-dependent dTAG^V^-1-mediated degradation of RPB1 in the mGFP-3XG linker RPB1 degron cell line using an antibody against the N-terminus of RPB1 and an anti-HA tag antibody. dTAG^V^-1 concentrations are the same as in Figure 1D. **(C)** Dose-response curve of RPB1 expression and dTAG^V^-1 concentration in the 6XG linker RPB1 degron cell line. **(D-F)** Flow cytometry analysis of RPB2 **(D)**, RPB3 **(E)**, and RPB4 **(F)** levels following treatment with different doses of dTAG^V^-1. Quantification of the median levels was performed for independent experiments. Bargraphs represent mean ±SEM. **(G)** Quantification of the median 5’-EU incorporation into nascent RNA following treatment with different doses of dTAG^V^-1 (see also Figure 1G). **(H)** Images showing 5’-EU incorporation into nascent RNA following 16 hours treatment with 500nM dTAG^V^-1. Scale bars=30um.

**Figure S3.**
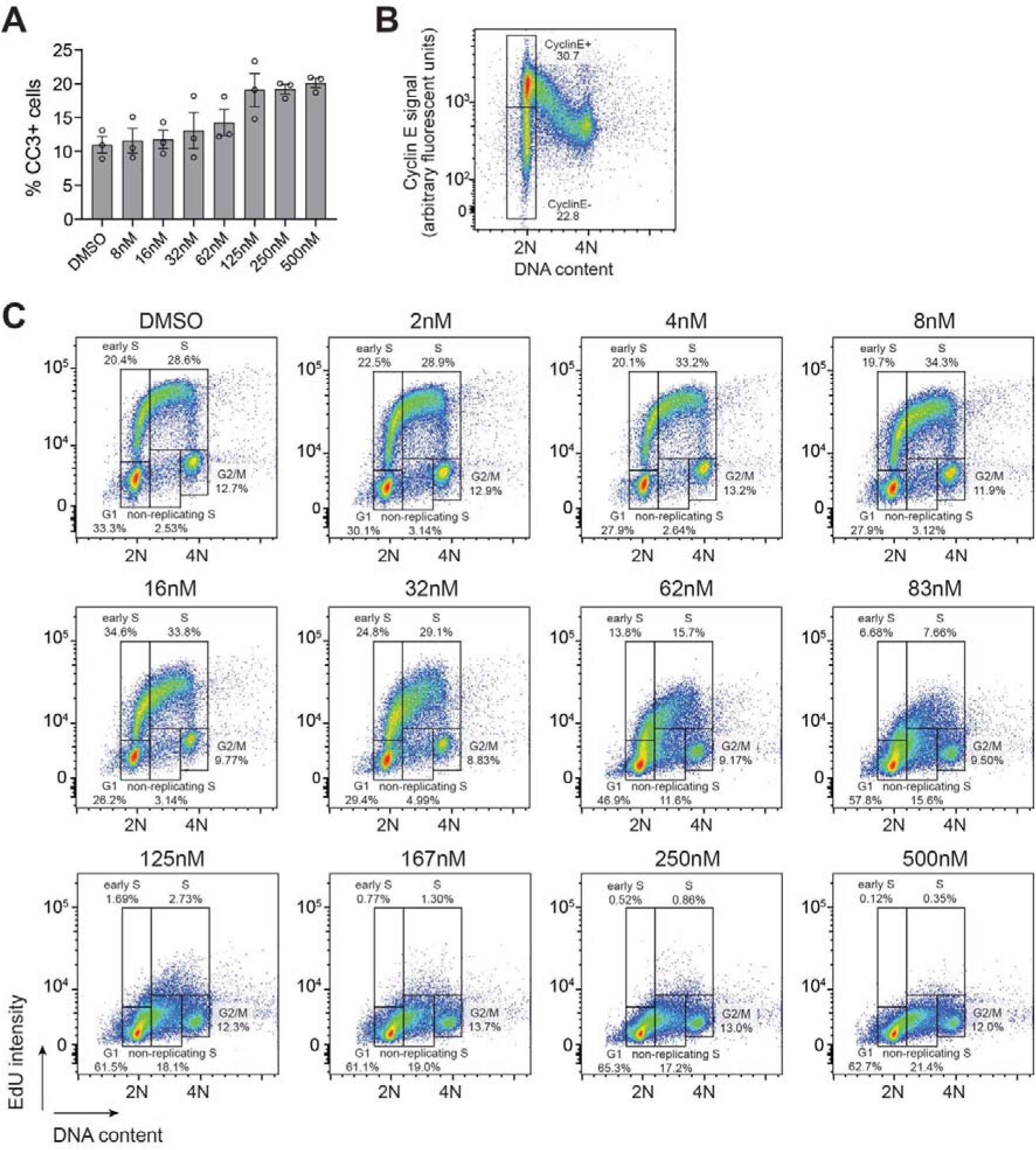
Related to. **Figure 1. (A)** Quantification of apoptotic cells, marked by cleaved caspase 3 (CC3) expression, using flow cytometry. Bargraphs represent mean ±SEM. **(B)** Dot plot showing a representative example of Cyclin E stain, which was used to mark cells in early and late G1 using flow cytometry. **(C)** Dot plots showing EdU incorporation following treatment with different concentrations of dTAG^V^-1. Representative gating strategy used for the quantification of Figure 1J is shown.

**Figure S4.**
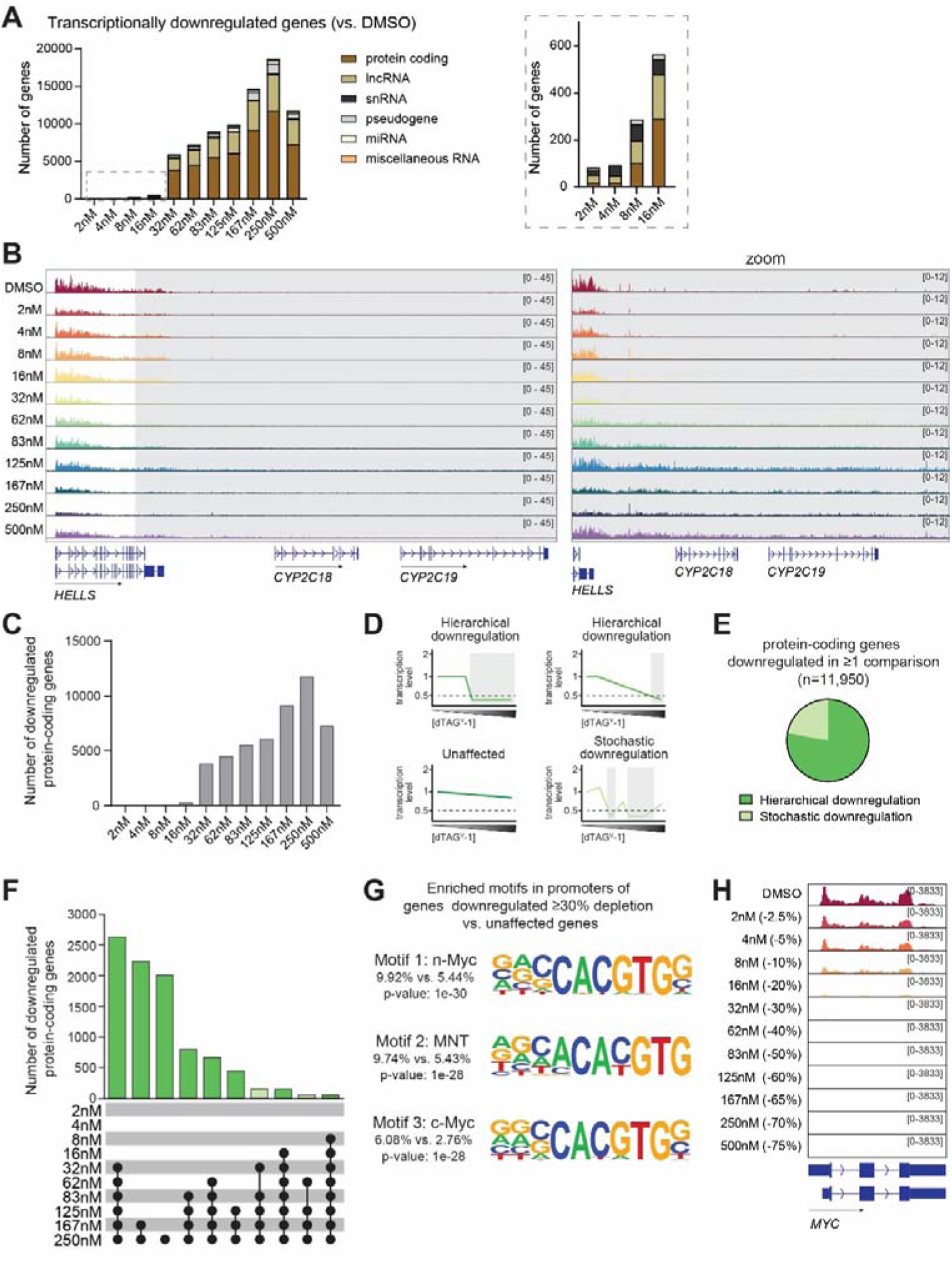
Related to. **Figure 2. (A)** Bar graphs with number of downregulated genes by biotype as determined by TT_chem_-seq. **(B)** Browser track with example of transcription readthrough. **(C)** Bar graphs with total number of downregulated protein-coding genes at each dTAG^V^-1 dose compared to DMSO. **(D)** Schematics showing examples of different transcription patterns across increasing dTAG^V^-1 concentrations. Genes whose transcription is consistently ≥2-fold downregulated (dotted line; grey background) from a specific dTAG^V^-1 concentration onwards, are binned as hierarchical. Genes whose transcription is never 2-fold downregulated at any dTAG^V^-1 dose are binned as unaffected. Genes whose transcription level are ≥2-fold downregulated at some dTAG^V^-1 concentrations, but do not meet the threshold consistently at higher dTAG^V^-1 concentrations are binned as stochastic. Color coding of the transcription pattern is consistent for (E) and (F). **(E)** Pie chart with the distribution of protein-coding genes expressed above 1 TPM, whose transcription is downregulated in ≥1 dTAG^V^-1 vs DMSO comparison. **(F)** Upset plot representing the intersection of the protein-coding genes downregulated across the individual dTAGV-1 vs DMSO pairwise comparisons. Analysis included all protein-coding genes expressed above 1TPM that are downregulated in ≥1 dTAG^V^-1 concentration vs. DMSO comparison. Bar graphs representing genes downregulated in a hierarchical pattern are shown in dark green, whereas bar graphs representing genes downregulated stochastically are shown in light green. Color-coding of bars is the same as used in (D) and (E). **(G)** Top 3 significantly enriched motifs in the promoter of genes downregulated at ≥30% depletion compared to unaffected genes. **(H)** Browser track showing transcription of *MYC* following treatment with different concentrations of dTAG^V^-1.

**Figure S5.**
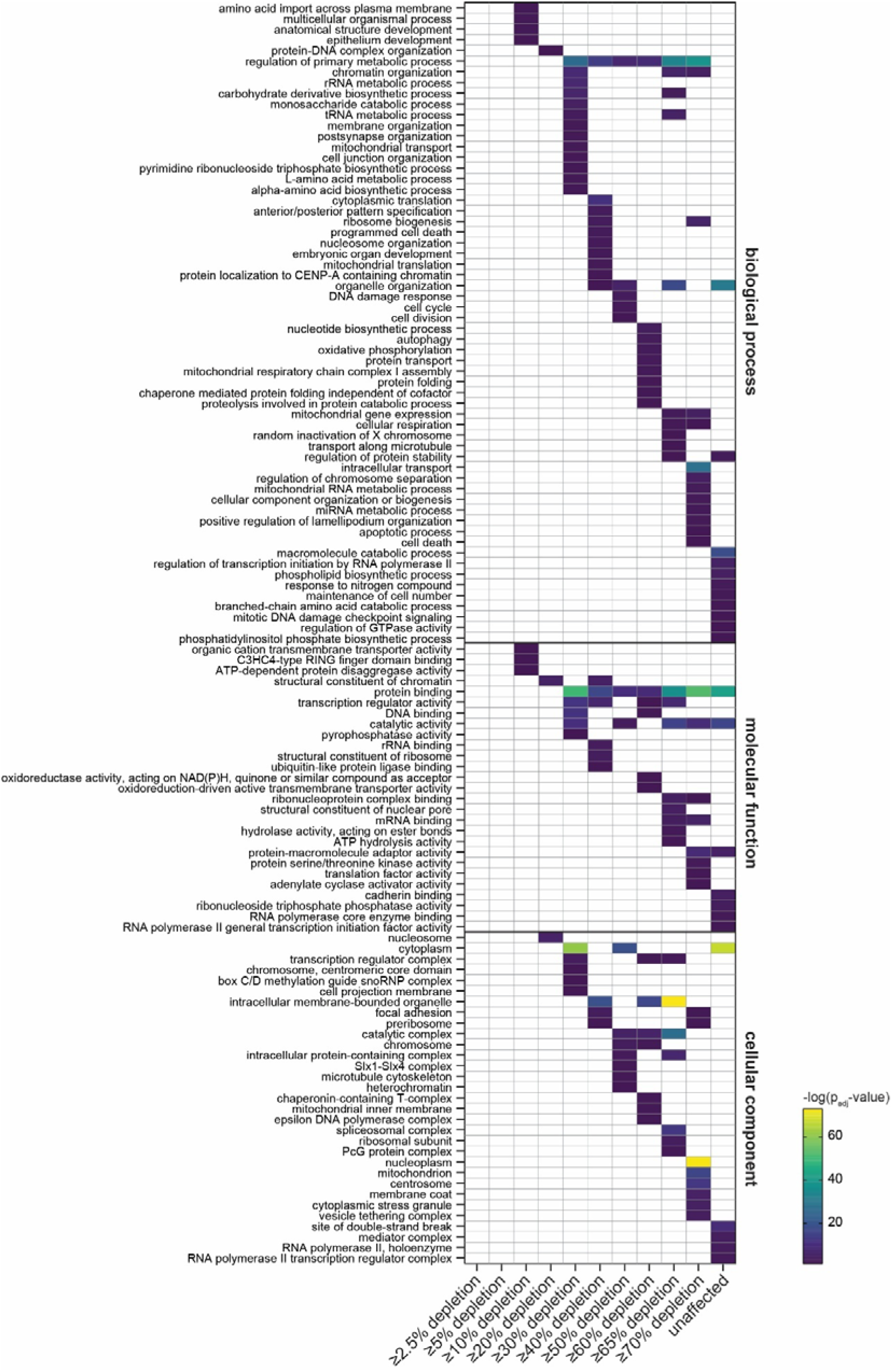
Related to. **Figure 2**. Heatmap with all gene ontology driver terms enriched for by genes downregulated in a hierarchical manner at each of the RPB1 depletion levels.

**Figure S6.**
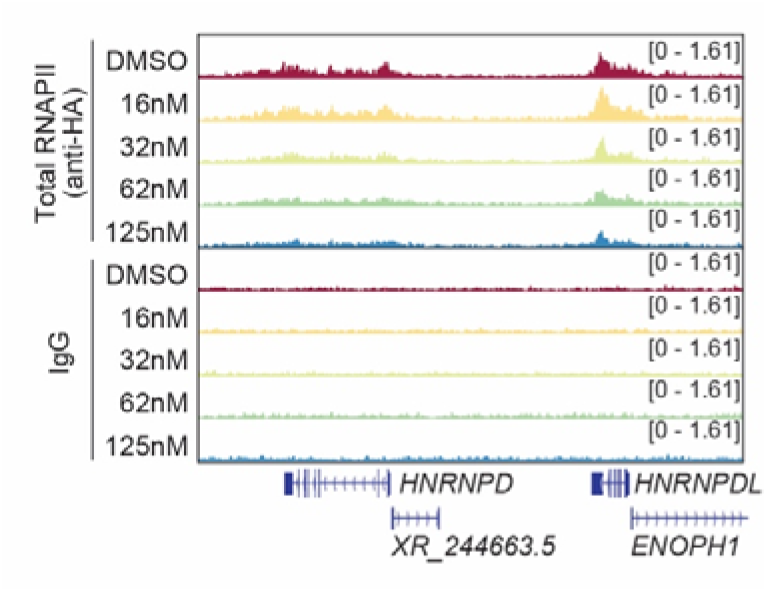
Related to. **Figure 3**. Browser track showing a representative example of CUT&RUN data for total RNAPII and IgG following treatment with different concentrations of dTAG^V^-1.

**Figure S7.**
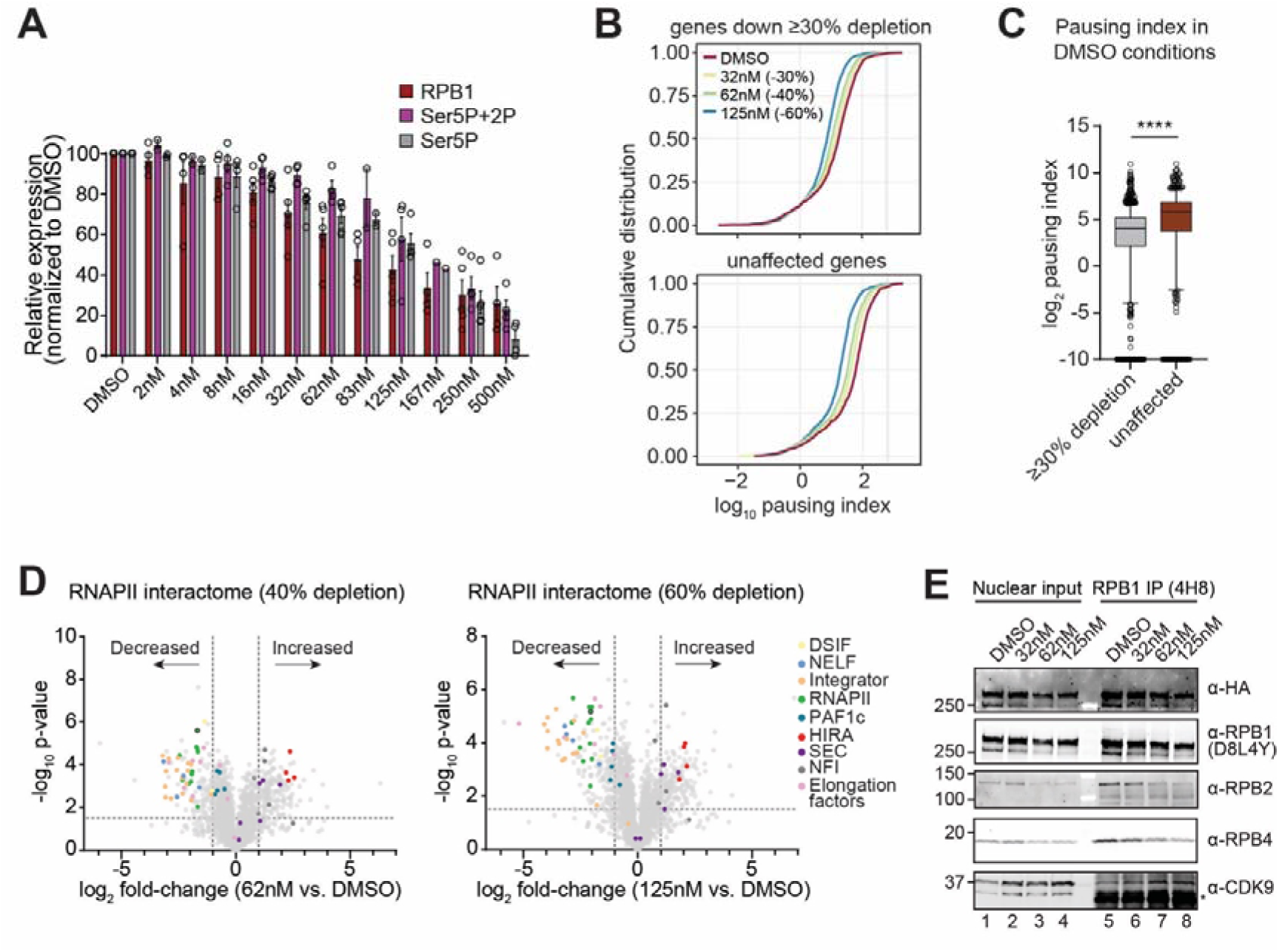
Related to. **Figure 4. (A)** Quantification of flow cytometry analysis of serine 5 and serine 5 + serine 2 phosphorylation on CTD of RPB1. Bargraphs represent mean of independent experiments ±SEM. **(B)** Empirical cumulative distribution function (ECDF) plots of pausing index for genes downregulated at ≥30% depletion and unaffected genes, following treatment with different concentrations of dTAG^V^-1. **(C)** Pausing index of genes downregulated at ≥30% depletion and unaffected genes in DMSO conditions. Statistical significance was determined using a Kolmogorov-Smirnov test. **(D)** The change in RNAPII interactome following 40% (left) and 60% (right) depletion of RPB1. Factors enriched following RPB1 depletion are shown on the right side of each volcano plot, while factors that are depleted from RNAPII following RPB1 depletion are shown on the left side of each volcano plot. **(E)** Western blot analysis of RNAPII subunits and CDK9 following immunoprecipitation of elongating RNAPII from nuclei of cells treated with different concentrations of dTAG^V^-1.

**Figure S8.**
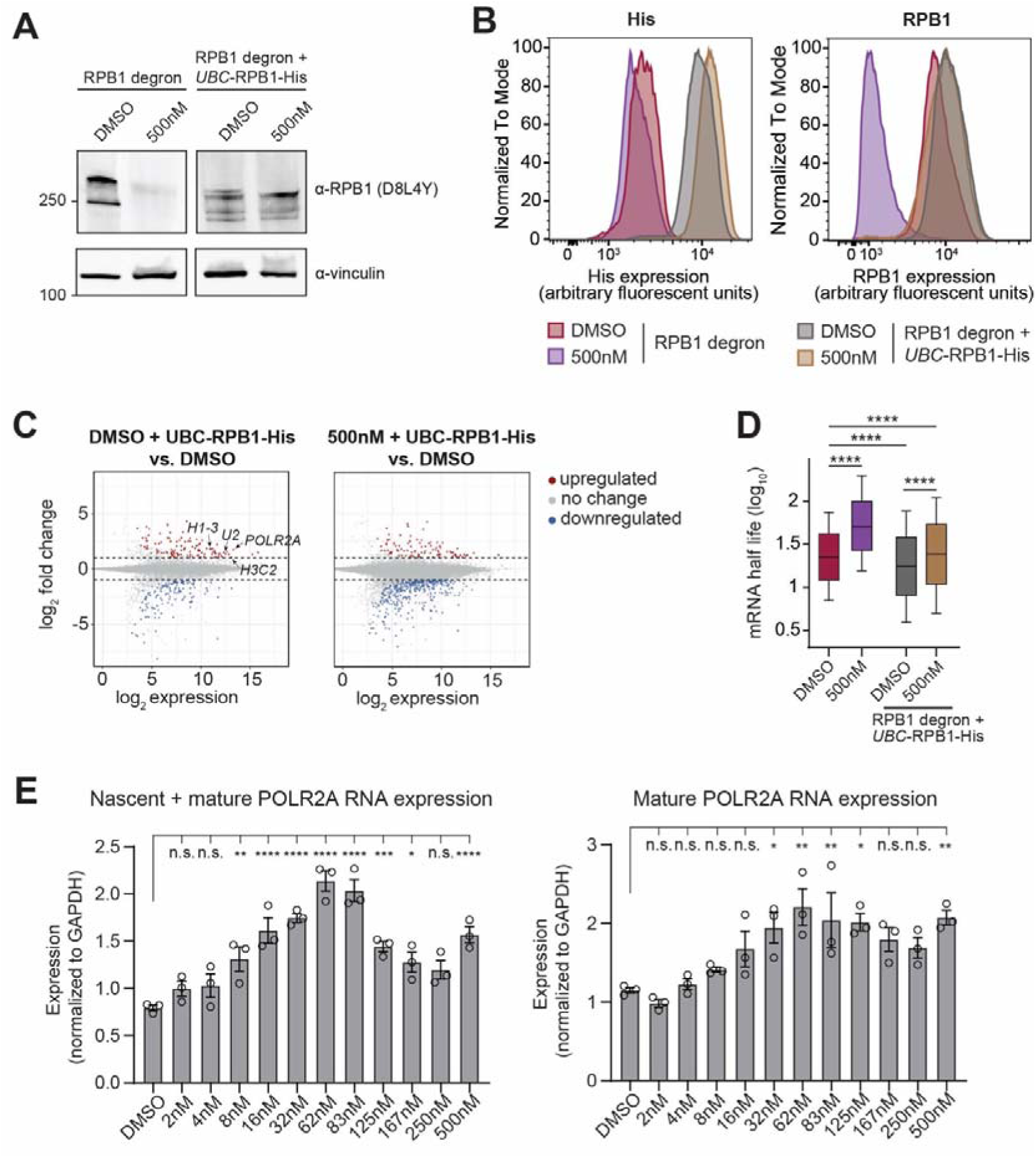
Related to. **Figure 7. (A)** Western blot analysis of total RPB1 expression following expression of the RPB1 transgene from a *UBC*-promoter and treatment with 500nM dTAG^V^-1. RPB1 expression was determined using an antibody against the N-terminus of RPB1. **(B)** Flow cytometry analysis of the RPB1 transgene (left) and total RPB1 (right) expression following expression of the RPB1 transgene from a *UBC*-promoter and treatment with 500nM dTAG^V^-1. RPB1 transgene expression was determined using an antibody against the His-tag, total RPB1 expression was determined using an antibody against the N-terminus of RPB1. **(C)** Scatter plots showing the differentially expressed genes in the TT_chem_-seq data of the RPB1 transgene-expressing cell line compared to the 6XG RPB1 degron cell line. Expression is basemean across all conditions, log2 fold-change has been shrunken using the apeglm method. Data are normalized to yeast spike-in. **(D)** Calculation of mRNA half-life expression of the RPB1 transgene. Data is shown as boxplots, with whiskers representing 5^th^-95^th^ percentiles. Statstical significance was determined using a Friedman test, followed by Dunn’s multiple comparisons test. **(E)** qRT-PCR results for the relative expression of *POLR2A* following treatment with different concentrations of dTAG^V^-1. Expression for total *POLR2A* RNA (left) and *POLR2A* mRNA (right) are normalized to *GAPDH*.

