## Supplementary Table 1 for "Systems-level feedback loops maintain gene expression homeostasis following RNA polymerase II dosage perturbation"

| **Name** | **Sequence** |
| --- | --- |
| **CRISPR-Cas9 gRNA** | |
| POLR2A_sgRNA_2_F | caccgccacccccgtgcatggcgg |
| POLR2A_sgRNA_2_R | aaacccgccatgcacgggggtggc |
| **Primers for plasmid construction** | |
| POLR2A_GIBS_F_sg2 | gttccgcgttacatagcatcgtacgcgtacgtgtttgggcgcagcgcgcctgcctccgccatgaccgagtacaagcccacggtgc |
| POLR2A_GIBS_R_sg2 | cagcattctagagcatcgtacgcgtacgtgtttggggggccacccccgtgcatggaagatccgccgccacccgacccaccac |
| NOE36_Backbone_F | tcttccatgcacgggggtggc |
| NOE36_BACKBONE_R | accttccagttttagaagctccacatcg |
| NOE34_2_F_Linker | tgtggagcttctaaaactggaaggtggc |
| NOE36_2_LINKER_R | gccacccccgtgcatggaagatccgccg |
| NOE36_B_F_Backbone | gcggcggatcttccatgcacggg |
| NOE36_B_R_Backbone | tttagaagctccacatcgaagacgaga |
| NOE36_B_F_15x | cactctcgtcttcgatgtggagcttctaaaactggaaggt |
| NOE36_B_R_15x | ccgtgcatggaagatccgccgccac |
| NOE37_3_backbone_F | agctgtacaagggtggcggtggctcgggcggt |
| NOE37_Backbone_R | ccttccagttttagaagctccacatcg |
| NOE37_2_F_mEGFP | ttcgatgtggagcttctaaaactggaaggtggcggtggctcgggcggtggtgggtcggtgagcaagggcgaggagc |
| NOE37_mEGFP_R | ccaccgccacccttgtacagctcgtccatgccg |
| UBC_promoter_gibson.F | atgtacgggccagatatacagaaacaggaagaagaacacattccc |
| UBC_promoter_gibson.R | cacccccgtgcatgctgtctaacaaaaaagccaaaaacggccag |
| **PCR primers** | |
| POLR2A.E10.F | tcacagcagtgcgcaaattc |
| POLR2A.E11.R | gagaagatttgcttgcctgtcc |
| GAPDH.F | cagcctcaagatcatcagca |
| GAPDH.R | tgagtccttccacgatacca |
| POLR2A.5UTR.F | agagacaaactgccgtaacctc |
| POLR2A.5UTR.R | tgggggaggaagaagaaaaagg |
| POLR2A.transgene.F | ccagccccacctacagtctc |
| POLR2A.transgene.R | tcgaggctgatcagcgggtttaa |
| TBX3.E2a.F | atgtacattcacccggacagc |
| TBX3.E2a.R | gggctggtatttgtgcatgga |
| POLR2A_N_seq02_F (genotyping) | tttccggtaagggaaagaaggg |
| POLR2A_N_seq02_R (genotyping) | gactcaggactccgaactgg |
| **HCR FISH probes** | |
| Scrambled-101.1 | gtccctgcctctatatctttatagggcctggaaacatcgcttcaa |
| Scrambled-101.2 | ggcaggatagtgtgcaagtagcgcattccactcaactttaacccg |
| POLR2A-58.1 | gtccctgcctctatatcttttcacactaagattcaagcgaaaggt |
| POLR2A-58.2 | cgtcaaagtctgcattgtacggagtttccactcaactttaacccg |
| POLR2A-50.1 | gtccctgcctctatatctttgaggagccagttgttaatgacagtc |
| POLR2A-50.2 | cccaatgccaatagtatgaccctcgttccactcaactttaacccg |
| POLR2A-40.1 | gtccctgcctctatatctttgcgtcgtacttcaccatcactgact |
| POLR2A-40.2 | acctggttgatggagttccgcacagttccactcaactttaacccg |
| POLR2A-23.1 | gtccctgcctctatatcttttgtacaccttgctgatctgctcgat |
| POLR2A-23.2 | tcttgttgtctgtctgtggcaagtgttccactcaactttaacccg |
| POLR2A-3.1 | gtccctgcctctatatctttgccagtgagttctaacagcacaagt |
| POLR2A-3.2 | gtggggtaggagtgagaacaccactttccactcaactttaacccg |
| oligodT-1.1 | gtccctgcctctatatctttttttttttttttttttttttttttt |
| oligodT-1.2 | tttttttttttttttttttttttttttccactcaactttaacccg |
